# Truncated radial glia as a common precursor in the late corticogenesis of gyrencephalic mammals

**DOI:** 10.1101/2022.05.05.490846

**Authors:** Merve Bilgic, Quan Wu, Taeko Suetsugu, Atsunori Shitamukai, Yuji Tsunekawa, Tomomi Shimogori, Mitsutaka Kadota, Osamu Nishimura, Shigehiro Kuraku, Hiroshi Kiyonari, Fumio Matsuzaki

## Abstract

The diversity of neural stem cells is a hallmark of the cerebral cortex development in gyrencephalic mammals, such as Primates and Carnivora. Among them, ferrets are a good model for mechanistic studies. However, information on their neural progenitor cells (NPC), termed radial glia (RG), is limited. Here, we surveyed the temporal series of single-cell transcriptomes of progenitors regarding ferret corticogenesis and, found a conserved diversity and temporal trajectory between human and ferret NPC, despite the large timescale difference. We found truncated RG (tRG) in ferret cortical development, a progenitor subtype previously described in humans. The combination of *in silico* and *in vivo* analyses identified that tRG differentiate into both ependymal and astrogenic cells. Via transcriptomic comparison, we predict that this is also the case in humans. Our findings suggest that tRG plays a role in the formation of adult ventricles, thereby providing the architectural bases for brain expansion.

## INTRODUCTION

A vast diversity of neurons and glia form functional neural circuits during the development of the cerebral cortex in mammals. These cells are progressively generated from multipotent neural stem cells, termed radial glia (RG), following genetic processes common across species, from initially neurons of the deep layers (DL), then neurons of the upper layers (UL), and finally glial cells, astrocytes, or oligodendrocytes (**Fig. S1A, S1B**; Rowitch and Kriegstein, 2010). These processes are also spatially organized with RG divisions in the ventricular zone (VZ), IPC-neuron differentiation in the sub-VZ (SVZ), and neuron migration to the cortical plate in an inside-out manner. In many mammalian phylogenic states, cerebral cortex evolved to gain an additional germinal layer (outer SVZ, OSVZ; Smart et al, 2002), where extensive neurogenesis and gliogenesis of outer RG (oRG) occur (Hansen *et al*, 2010; Fietz *et al*, 2010; Reillo *et al*, 2011; Gertz & Kriegstein, 2015) resulting in the amplification of neuronal and glial populations in the cortex (**Fig. S1**; Rash et al, 2019). On the other hand, a new subtype of NPC in the VZ has recently been reported in humans and rhesus macaques (deAzevedo et al, 2003; Nowakowski et al, 2016; Sidman and Rakic, 1973), lacking the basal attachment and is therefore termed truncated RG (tRG). However, how widely tRG appears in gyrencephalic (or even lissencephalic) mammal development, what mechanisms underlie their formation, and what they produce remain unknown. As distinct NPC subtypes may possess different capacities to generate differentiated progenies (Huang et al, 2020; Rash et al, 2019), it would be critical to find answers to those questions to understand commonality and diversity in the mammalian brain evolution.

Genetic manipulation of individual cell types *in vivo* and single-cell transcriptome analysis are two major and successful approaches in revealing the properties of cells, such as their proliferation and differentiation. Thus, single-cell RNA sequencing (scRNA-seq) of the human brain during development has been extensively performed (Herring *et al*, 2022; Bhaduri et al, 2020, 2021; Huang et al, 2020; Johnson et al, 2015; Liu et al, 2017; Polioudakis et al, 2019; Pollen et al, 2015; Zhong et al, 2018). However, the *in vivo* behavior in humans and other primates and the underlying mechanisms remain less explored owing to the limited experimental access to the developing primate cortices. Particularly, resources available for the late human embryonic brain are extremely rare due to ethical challenges. Also, studies using brain organoids face issues in recapitulating the specification and maturation of cell types during human brain development (Bhaduri et al, 2020). In this context, the ferret (*Mustela putorius furo*) is highlighted as a suitable animal model, which compensates for the difficulties in studying human cortical development. Ferrets are carnivores that develop common gyrencephalic features, such as the OSVZ and a folded brain, and are frequently used in mammalian models of brain development and circuit formation because neurogenesis continues in their early neonatal stages (Chapman & Stryker, 1992; McConnell, 1988; Noctor *et al*, 1997; Jackson *et al*, 1989). Since this species is experimentally manipulable, recent studies have developed *in vivo* gene manipulation and editing technology using *in utero* electroporation (IUE; Matsui et al, 2013; Kawasaki et al, 2012; Tsunekawa et al, 2016; timelapse imaging; McConnell, 1995; Chenn & McConnell, 1995). Furthermore, ferrets showed severe microcephalic phenotypes via *ASPM* (*Abnormal Spindle-like Microcephaly-associated* gene) knockout (Johnson et al, 2018; Kou et al, 2015), which greatly differs from a minor phenotype in mouse *ASPM* knockout mutants (Capecchi and Pozner, 2015; Fujimori et al, 2014; Jayaraman et al, 2016; Pulvers et al, 2010). This remarkable finding suggests the presence of mechanisms underlying brain enlargement and circuit complexity shared by gyrencephalic species to some extent. Results from transcriptome profiling of ferret cortical cells has revealed regional differences in germinal layers and cell-type composition (Johnson et al, 2018; de Juan Romero et al, 2015). However, because of the incomplete genomic information, especially due to the lack of genetic models, the databases with ferret data have been less reliable than those with human or mice data. This has posed a limit for the accurate comparison of single cell transcriptomes between ferrets and humans. Hence, the temporal pattern of molecular signatures of ferret NPC remains largely unexplored at single-cell resolution. Comparison of progenitor subtypes and sequential events at the single-cell transcriptome level regarding development between ferrets and humans will greatly help to recognize common and species-specific mechanisms underlying the construction of a complex brain.

In this study, we comprehensively analyzed the developmental dynamics of progenitor populations during ferret corticogenesis to clarify shared or species-specific mechanisms of corticogenesis in gyrencephalic mammals and found that ferrets generate tRG at late neurogenic and early gliogenic stages like humans and other primates. Analysis of the pseudotime trajectory and temporal histochemical pattern suggested that tRG generate ependymal and astrogenic cell fates. Then, we compared temporal series of ferret and human single cell transcriptomes (Bhaduri et al, 2021; Nowakowski et al, 2017). Remarkably, we found homologous temporal progenitor trajectories irrespective of the large differential corticogenesis timescale. This study combining rich ferret and human transcriptome data with *in vivo* analysis using ferrets emphasizes the value of single-cell ferret transcriptome datasets.

## RESULTS

### Temporal patterns of neurogenesis and gliogenesis in the cerebral cortex of ferrets

First, we confirmed histochemically the spatial and temporal pattern of RG, oRG, and IPC and the appearance of diverse cortical neurons to determine the period in which samples were to be taken for scRNA-seq (**Fig. S1**). While neurogenesis mostly terminated by P5 in the dorsal cortex (**Fig. S1A, S1B**), oligodendrocyte progenitors of dorsal origin (OLIG2^+^) and RG with gliogenic potential (GFAP^+^) became detectable approximately at E40 and increased in number postnatally (**Fig. S1C, S1D**). To recapitulate neurogenesis and gliogenesis in the ferret cortex at a high resolution, we decided to determine the temporal trajectories of cell types in the course of ferret brain development (schematically represented in **Fig. 1A**) by performing single cell transcriptomes of neural progenitors, neurons and glial cells. Based on the above histochemical observations (**Fig. S1A, S1B**), we examined single cell transcriptomes at the six developmental time points: embryonic days E25, E34, and E40 and postnatal days P5 and P10. We carried out single-cell RNA-sequencing (scRNA-seq) of ferret dorsal cortex at these six developmental time points (**Fig. 1**). We prepared two series of cell populations isolated in different ways to enrich the progenitor subtypes (**Fig. 1B**): (1) FACS-based sorting of the neural stem cell fraction labeled with an AzamiGreen (AG)-driven *HES5* promoter (Ohtsuka et al, 2006) and (2) collecting cells forming the VZ, SVZ, and intermediate zones (IZ) of cerebral cortices after discarding the cortical plate (CP, **Fig. 1B, S2A**).

**Figure 1.**
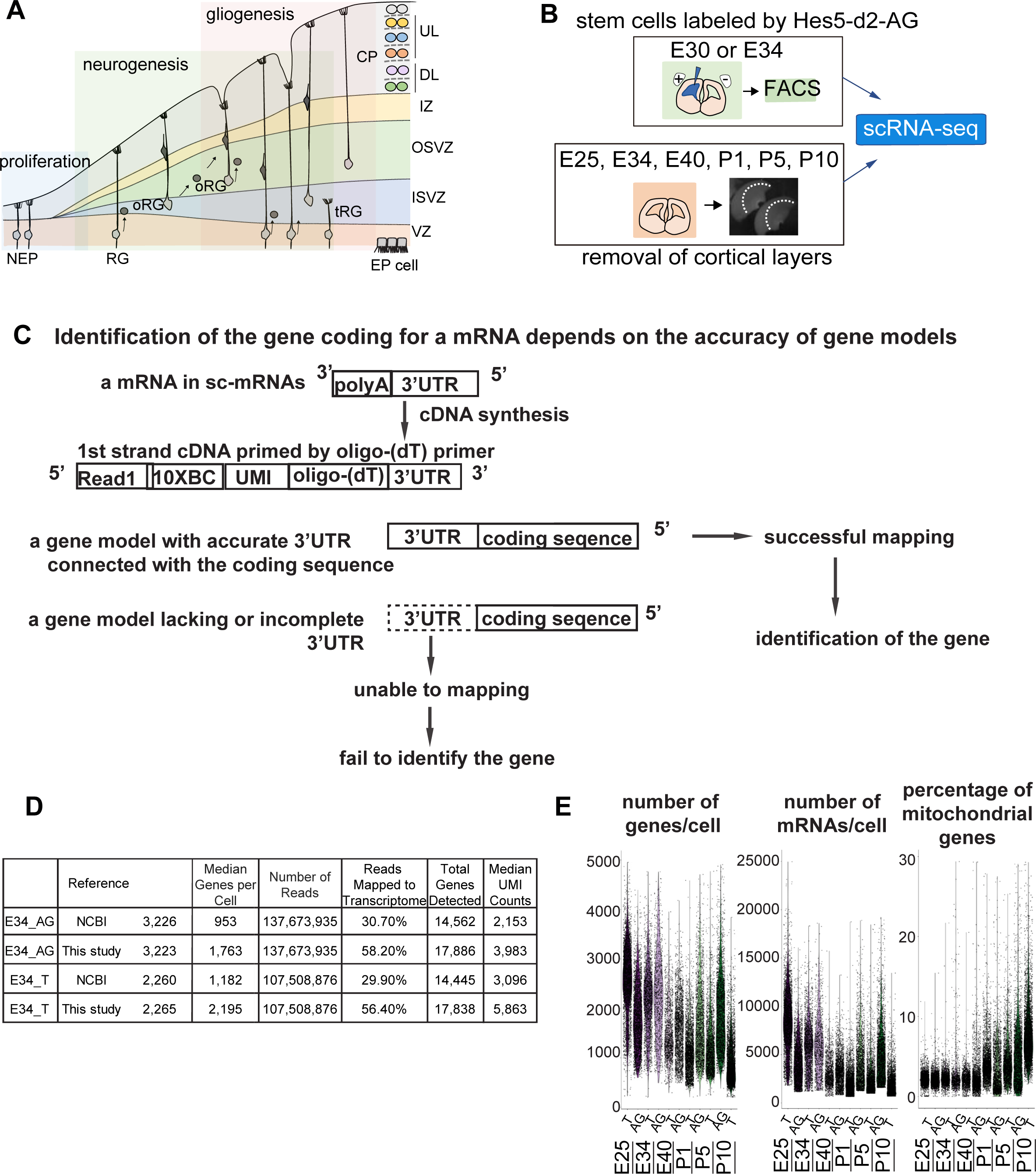
Improvement of the gene model for scRNA-seq of ferretes. **A** Schematic representing cortical development and emergence of diverse neural progenitors in humans and ferrets. Progenitors sequentially generate DL- and subsequent UL-neurons, and finally glial cells, and form ependymal cells. RG and oRG have been morphologically and positionally identified in both humans and ferrets while tRG have been only reported in humans. VZ, ventricular zone; ISVZ, inner subventricular zone; OSVZ, outer subventricular zone; IZ, intermediate zone; DL, deep layer; UL, upper layer; CP, cortical plate; NEP cell, neuroepithelial cell; Mn, migrating neuron; and EP cell, ependymal cell. **B** Schematic representing the experimental design and time points used to build the transcriptome atlas of developing somatosensory cortex of ferrets. Single cells were isolated using 10X Chromium (see **Materials and Methods**). **C** Linkage of the 3’UTR sequence and the coding sequence of genes has been improved in this study. This linkage is necessary to assign the gene coding for a mRNA as far as 10x Genomics Chromium kit is used for scRNA-seq (see **Materials and Methods**). **D** Table comparing quality-control metrics of an alignment with either MusPutFur 1.0 (UCSC gene models) or MusPutFur 2.60 (this study) using E34 samples. The total number of genes detected and median genes per cell were higher with MusPutFur 2.60. **E** Violin plots showing the number of genes, mRNAs and the percentage of mitochondrial genes per cell in each sample and time point.

### Improvement of the gene model for scRNA-seq of ferrets

High quality information about gene models over the entire genome is prerequisite to obtain a high resolution of single cell transcriptomes from scRNA-seq. Despite several previous transcriptome studies on the ferret cortex (Johnson et al, 2018; de Juan Romero et al, 2015), public information on the ferret genome is incomplete; identification of the gene corresponding to a cDNA requires its accurate 3’-untranslated region (3’-UTR) and its correct connection to the coding sequence, because single cell-cDNA library were made by Oligo (dT)-priming method (10x Genomics Chromium, **Fig 1C)**. However, information regarding the 3′-UTR of genes had been poor for ferret genomic datasets, which has impeded high resolution single-cell transcriptome analyses that can offer an accurate comparison with human datasets, Thus, we first improved the annotations of ferret genomic DNA sequences using Chromium droplet sequencing, which tagged all contigs from a long genomic DNA in a droplet, and constructed new gene models based on the improved genomic annotations and newly obtained RNA-seq reads of various tissue types (**Table S7 and see Materials and Methods**). Comparing with NCBI references, mapping scRNA-seq reads from E34 to our own references not only improved the mapping rate (number of reads mapped to the transcriptome) from ∼30% to ∼56%, but also increased the median number of genes detected from ∼1000 to ∼2000. (**Fig. 1D**). We found similar mapping rate and median number of genes detected across all sampling stages (**Table S1**).

### Single-cell RNA-seq revealed subtypes of cortical cells in ferrets

We then combined scRNA-seq information from the two cell populations on all the sampling stages for unbiased clustering. **Fig. 2A** shows cell clustering projected in the Uniform Manifold Approximation and Projection (UMAP) space (preprint: McInnes et al, 2018; Stuart et al, 2019). We characterized 26 transcriptionally distinct clusters from 30,234 ferret cortical cells through corticogenesis and detected up to 2,600 median genes per cell (**Fig. 1D, E; Table S1**). After combined clustering of the two collectives of cell populations prepared by independent methods, we re-separated cells into each collective, and confirmed a reproducibility of clustering between two different collectives (**Fig. S2B**).

**Figure 2.**
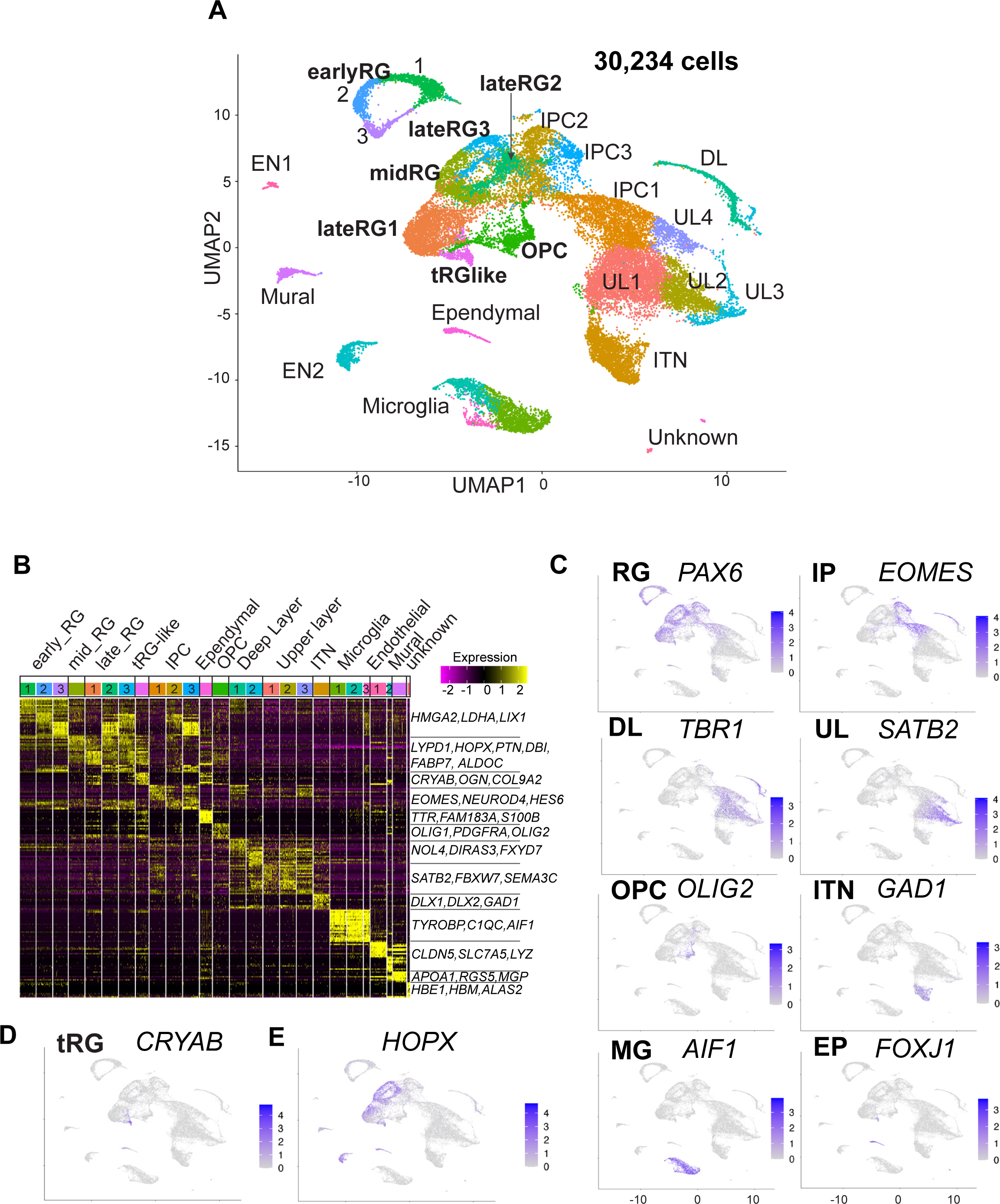
Single-cell RNA-seq reveals ferret transcriptome signatures and the cell types. **A** UMAP visualization showing cells colored by Seurat clusters and annotated by cell types (as shown in **C, D**). **B** Heatmap showing expression profiles of cluster marker genes based on log fold-change values. Cells were grouped by Seurat clustering (transverse). Cell types were assigned according to the expression of marker and differentially expressed genes in each cluster. Early RG was defined by the expression of *HMGA2, LDHA,* and *LIX1*; and late RG, by *PTN, ALDOC,* and *FABP7*. OPC, oligodendrocyte precursor cell; ITN, interneuron; and EN, endothelial cells. Other cell type abbreviations are shown in Fig. 1A and the main text. Color bar matches the Seurat clusters in (**A**). The ten most enriched representative genes in each cluster are shown, with typical marker genes noted in (**C**–**E**). **C** Normalized expression levels of representative marker genes of different cell types projected onto the UMAP plot in (**A**). MG, microglia. **D** Expression pattern of *CRYAB*, a marker for tRG, only described in humans and other primates. **E** The expression pattern of *HOPX*, a marker for oRG in humans and other primates in the UMAP plot.

We also examined whether cells with high mitochondrial contents affect clustering of ferret single cells, because we have decided to include all cells that had less than 30% of mitochondrial genes in our analysis based on the percentage of reads mapped on the mitochondrial genome; we found that the majority of cells in each cell type had a value less than 5% while some cells contained them in the range between 0% and 10%, up to a maximum of 28% (**Fig, 1E**). We confirmed that single cells after filtering cells with the threshold of 10% mitochondrial content (28,686 cells in our dataset) and those with the threshold of 30% (**Fig. 2A**), provided similar clustering patterns with each other, both of which 26 clusters **(Fig. S2C)** indicating that the current selection of cells with the mitochondrial contents is appropriate.

Cell clusters were annotated according to their specific gene expression patterns (**Fig. 2B, C, S3A, Table S1**) and assigned into ten cell types: RG (early, mid, and late), IPC, OPC, ependymal cells, excitatory cortical neurons (DL and UL), inhibitory neurons (ITN), microglia, endothelial cells, mural cells, and unknown cells (**Fig. 2A**). RG, IPC, and neuronal clusters were aligned according to the neuronal differentiation process in the UMAP plot (**Fig. 2A**). RG cells were classified into three clusters (early, mid, and late) according to their collection stages and also the expression of temporally altered RG markers reported previously (hereafter named temporal markers) (Okamoto et al, 2016; Telley *et al*, 2019). The “early RG” clusters comprised E25 cells while the “mid RG” clusters, mostly E34 cells (**Fig. S3B**). Early and late RG, and IPC were subdivided into three subclusters that expressed different cell cycle markers (**Fig. S3C**).

Remarkably, we identified a small cluster (409 cells) of *PAX6*-expressing RG subtype, 69% of which expressed *CRYAB* (**Fig. 2D**). *CRYAB*, encoding a molecular chaperone (Yamamoto et al, 2014) is a unique marker for human tRG (Nowakowski et al, 2016); therefore, we designated them as tRG-like cells in ferrets. In contrast, oRG cells failed to be distinguished from ventricular RG cells (vRG) in ferrets by unbiased clustering alone (see below). HOPX, a typical marker for oRG in human tissues (**Fig. 2E, S3D**; Pollen et al, 2015), has been indeed detected in both oRG and vRG at late stages *in vivo* in ferrets (**Fig. S3D**; Johnson et al, 2018; Kawaue et al, 2019).

### tRG emerge around birth during the development of somatosensory cortex in ferrets

The presence of a tRG-like cluster in ferret transcriptomes led us to investigate the properties of the cells in this cluster. To examine their cell morphology, we sparsely labeled NPC by electroporating P0 embryos with an expression vector for enhanced green fluorescent protein (EGFP) and found a cell population showing major characteristics of human tRG cells (Nowakowski et al, 2016): expression of CRYAB and PAX6 with an apical endfoot and truncated basal fiber in the VZ during late neurogenesis (**Fig. 3A,B, Supplementary movie1, 2**). CRYAB expression emerged shortly prior to birth (day 41), gradually increasing in cell number, and became mostly restricted to tRG-shaped cells in the VZ and SVZ (**Fig. 3C, 3D, S4A**). These CRYAB^+^ cells were mostly post-mitotic (KI67^-^) and neither IPC (TBR2^+^) nor OPC (OLIG2^+^) at P10, while a small fraction of those cells expressed these markers at P5 (**Fig. 3E–3G**). These histochemical observations were consistent with our single-cell transcriptome data (**Fig. S4A–F**). Altogether, we concluded that tRG-like cells in ferrets are equivalent to human tRG cells (as also confirmed by transcriptomic comparison between humans and ferrets, shown later). A difference in the late neurogenic cortex between humans and ferrets is that the VZ and OSVZ are connected by conventional RG cells in ferrets (**Fig. 3A, S4G**), but separated in humans due to the disappearance of conventional RG cells (Nowakowski et al, 2016).

**Figure 3.**
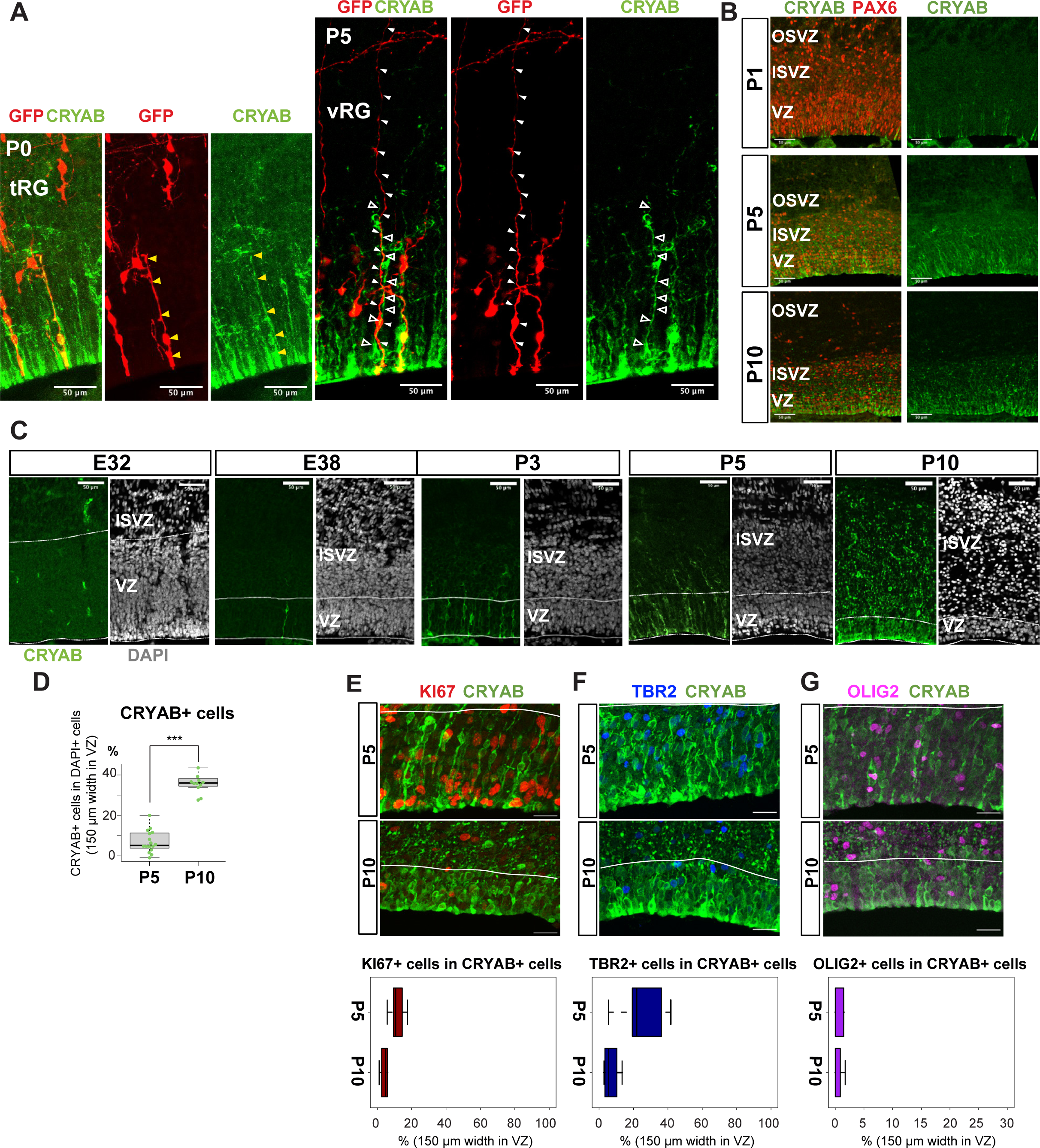
Truncated tRG emerges during postnatal cortical development in ferrets. **A** Representative images showing the cellular features of tRG and aRG in VZ at P0, and P5, respectively, stained for GFP (red) and CRYAB (green). RG cells were sparsely labelled with a GFP-expressing plasmid at E30 via IUE for P0 samples, and at P3 for P5 samples. MAX projection was performed on a 30 µm vibratome section with 5 µm or 2.5 µm interval for each z-image at P0 or P5, respectively. Scale bar, 50 µm. **B** Expression patterns of CRYAB (green) and PAX6 (red) in ferret germinal zones during postnatal development (P1, P5, and P10) (cryosection thickness = 12 µm). **C** Developmental profile of CRYAB expression in ferret cortices. Immunostaining for CRYAB (green) and DAPI (gray) on cryosections at E32, E38, P3, P5, and P10 (cryosection thickness = 12 µm). **D** Quantification of CRYAB^+^ cells among all nuclei (DAPI) in VZ strips through immunostaining, indicating a great increase in CRYAB-expressing cells (strip width = 150 µm; n = 3 for P5; n = 2 for P10; Wilcoxon rank-sum test *p*-value = 6.574e-08). **E–G** Expression of other markers in CRYAB^+^ cells. Representative images taken with a 100X- objective are shown with MAX projection. The border of VZ is shown with a white line. Below each double staining image, quantification of KI67-, TBR2-, or OLIG2-expressing cells among the CRYAB^+^ cell population in the VZ is shown (n = 2 for P5; n = 2 for P10 for each staining except n = 3 for TBR2 staining). Box and whisker plots indicate ranges (lines) and upper and lower quartiles (box) with the median. Scale bars, 20 µm.

### Temporal fates of RG cells are predicted by pseudo-time trajectory analysis

To understand the relationship between various cortical progenitors in the developing ferret brain, particularly in the origin and fate of ferret tRG, pseudo-time trajectory analysis was performed. Single cells that had been subjected to single-cell transcriptome analyses (from E25 to P10) were unbiasedly ordered along a trajectory based on their transcriptome profiles (**Fig. S5A, S5B**). To simplify our analysis, we first excluded interneurons, microglia, endothelial cells, and excitatory neuronal clusters. Subsequently, 6,000 single cells were randomly selected from the remaining cell population for further analysis. The pseudo-time analysis predicted a reasonable trajectory consisting of three branching points that generated seven states (**Fig. 4A, S5B**). We assigned major cell types for each state based on the cell clusters defined by UMAP analysis (**Fig. 3A, Table S2**). The trajectory successfully predicted cortical development, along which *HES1*^+^ stem cells shift their features from the earlier to the later stages, generating branches of differentiated cells (**Fig. 4B, 4C, S5D**). Major *HES1*^+^stem cell states (state 1, 3, 5 and 7) contained cells at different stages; thus, termed NPC1, 2, 3 and astrogenic, respectively (**Fig. S5C**). NPC1 (mostly from E25) bifurcated into the neuronally differentiated lineage and NPC2 state (**Fig. S5B**), which produced the OPC lineage and NPC3 state at the second branching point. When neurogenesis gradually declined after birth, the NPC3 state, mainly consisting of late RG cells and tRG (**Fig. S5C**), adopted two states, state 6 with an presumptive ependymal fate (*FOXJ1*^+^, *SPARCL1*^+^) and state 7 with presumptive astrogenic fate (*FOXJ1*^-^*, SPARCL1*^+^), as judged by the combination of late fate markers (Liu et al, 2022; Saadoun and Papadopoulos, 2010; Yu et al, 2008, **Fig. S3A, 3D**).

**Figure 4.**
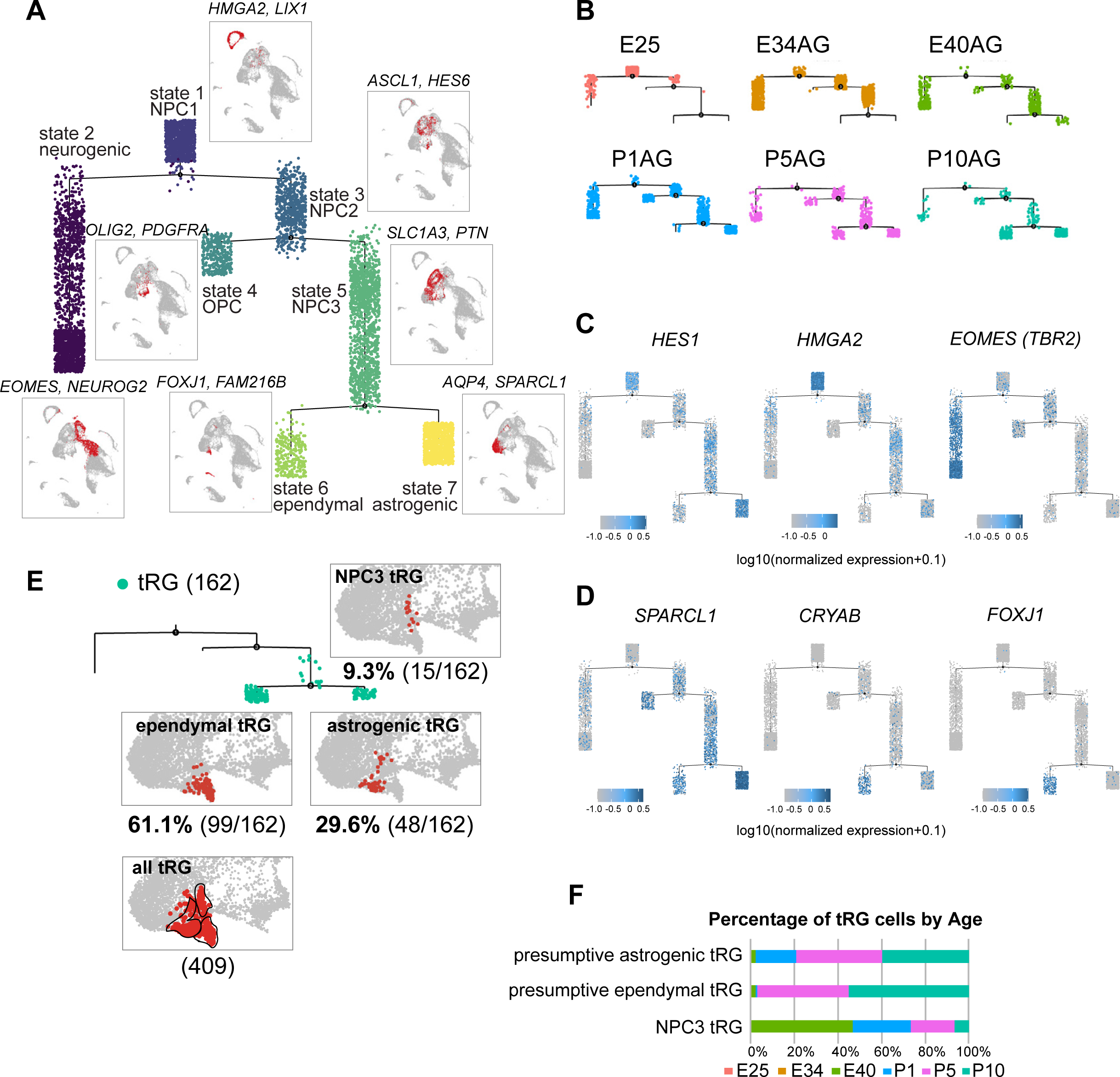
Temporal fates of RG cells predicted by pseudo-time trajectory analysis. **A** Pseudo-time trajectory tree of progenitor cells in the ferret developing cortex (assembled with Monocle v2). Cellular distribution at each state is the same as in the UMAP plot shown in Fig. 1C. Cell types representing each state and their gene markers are shown next to each state (see below). **B** Trajectory trees split by collection stages (AG samples are shown). **C** Distribution of cells expressing marker genes for stem cell states (*HES1* and *HMGA2*) and the neurogenic state (*EOMES*) along trajectories. Color densities indicate the log-normalized unique molecular identifier count for each gene. **D** Normalized expression of three genes: *SPARCL1, CRYAB,* and *FOXJ1* in the state 6 and state 7 cells in the pseudotime trajectory tree (A). The combination of expression levels of the four markers gene discriminates state 6 (ependymal) and state 7 (astrogenic). See the text for details. **E** Distribution of tRG cells along the tree and tRG-focused UMAP visualization. The tRG cells (n = 162) were found on the three states, NPC3 (9.3%; 15 cells), ependymal-tRG (61.1%; 99 cells), and astrogenic-tRG (29.6%; 48 cells). **F** Composition of the three types of tRG in (D) by the collection stage (AG and T samples are combined). No tRG cells were collected from E25 and E34.

Remarkably, tRG cell population was distributed into three branches along the pseudo-time trajectory (**Fig. 4D, 4F**): NPC3 (mainly during E40–P1), astrogenic (P1–P10), and ependymal (P5–P10) branches, suggesting that tRG prenatally arise as precursors of ependyma and astroglia in the ferret cortex (see below). Different territories were assigned into these three states in the UMAP even when we examined the entire tRG population (409 cells) before making a random selection of 6,000 cells for pseudotime trajectory analysis, supporting the reliability of this presumptive categorization of tRG (**Fig. 4E**).

### tRG cells is likely to possess ependymal and gliogenic potential during cortical development in ferrets

To test the hypothesis that tRG generate both ependymal cells and astroglia populations, we first examined the expression of an ependymal marker, FOXJ1, a master regulator of ciliogenesis, according to CRYAB expression in the VZ. We observed that CRYAB^+^ tRG cells gradually co-expressed FOXJ1, reaching up to 90% co-expression by P10 (**Fig. 5A, 5B**). Similarly, our transcriptome data showed that the fraction of *CRYAB-FOXJ1* double-positive cells increased in the tRG cluster from P1 to P10, whereas other RG clusters maintained low *CRYAB* expression (**Fig. 5C, 5D**). From P5 onwards, differentiating ependymal cells (*CRYAB-FOXJ1* double-positive) accumulated within the VZ (**Fig. 5A**). These cells often align their nuclei in parallel to the ventricular surface, near which nuclear lined aggregates are observed more frequently (between two arrowheads in P5 and P10 in **Fig. 5A**), and their cell body finally settled on the apical surface with short basal fibers by P14 (**Fig. 5E**). These data suggest that a majority of tRG cells progressively upregulates *FOXJ1* expression to adopt an ependymal cell fate during postnatal development. Concomitantly, adenine cyclase III expression in both primary and multiciliated cells indicated that ciliogenesis progressed postnatally, forming multi-ciliated ependymal cells on the ventricular surface of the ferret cortex by P35 (**Fig. S6A**). Altogether, it is most likely that FOXJ1^+^tRG are fated to be ependymal cells that constitute the ventricular surface. To prove this, it is necessary to chase tRG by knocking in a fluorescent marker gene via *in utero* electroporation, or to follow the change in cell shape from tRG to a characteristic ependymal morphology in slice cultures by labeling with a fluorescent gene via *in utero* electroporation. However, the latter would not be realistic, because the process of the transformation from tRG to ependymal cells is slow to take a long time in an order of 10 days. It is an important future issue to develop the strategy to genetically follow the fate of tRG.

**Figure 5.**
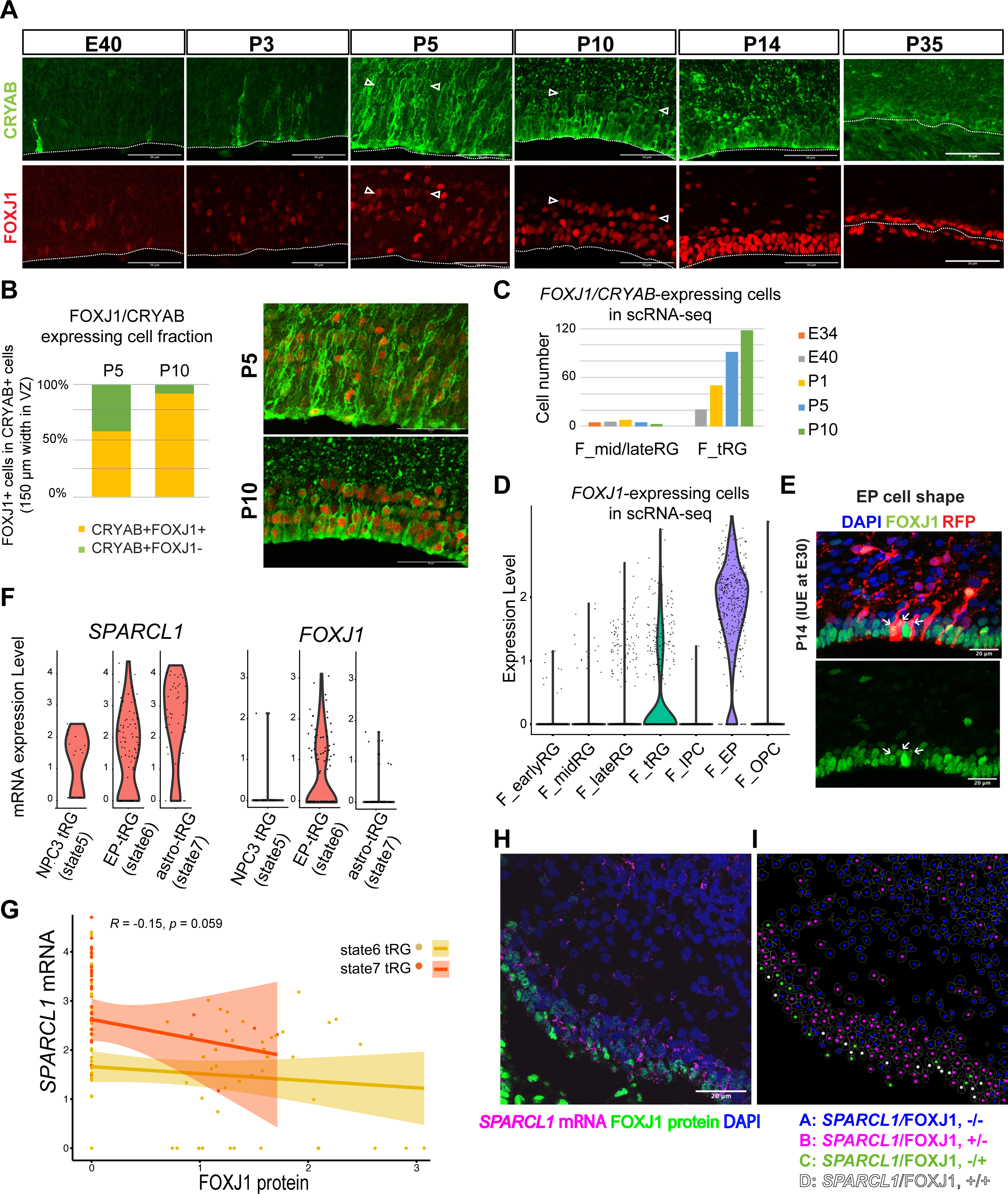
tRG adopt both ependymal cells and astrogenic fates in ferrets. **A** Immunostaining of 12 µm cortical cryosections with CRYAB (green) and FOXJ1 (red), focusing on the VZ at E40, P3, P5, P10, P14, and P35. Scale bar, 50 µm. CRYAB and FOXJ1 double positive nuclear rows are shown with arrowheads at P5 and P10 when this nuclear alignment is visible. **B** Percentage of FOXJ1 expression in CRYAB-expressing cells (n = 3 for P5; n = 2 for P10; Wilcoxon rank sum test *p*-value = 1.749e-05). Images with merged channels in A are shown with the same color codes, antibodies and scale bars as A. **C** Number of cells expressing both *CRYAB* and *FOXJ1* in the mid and late RG, or tRG clusters shown in Fig. 1C (with unique molecular identifier counts higher than 0.25). Colored bars indicate the stages of sample collection. *CRYAB*- and *FOXJ1*-expressing cells increased over time only in the tRG cluster. **D** Normalized expression levels of *FOXJ1* in each cell in the indicated ferret clusters in Fig. 1C. **E** Cortical origin and shape of FOXJ1-expressing ependymal cells indicated with white arrows. Staining for FOXJ1 at P14 after labeling cortical progenitors with an mCherry-expressing vector via IUE at E30. The maximum projection images with 1 µm z-step size are shown. Cryosection thickness = 12 µm; scale bar = 20 µm. **F** Normalized expression of *SPARCL1* and *FOXJ1* in each cell in ferret tRG clusters separated by the pseudo-time trajectory analysis (see **Table S3**). **G** Relationship between *SPARCL1* and *FOXJ1* transcripts in individual cells within each tRG cluster. Within the 15 cells that were classified in state 5 tRG, only one expressed *FOXJ1* mRNA. Pearson relationship analysis was performed using cells from state 6 and 7 (R=0.19 p value=0.018). **H** Immunostaining of a P10 cortical tissue for FOXJ1 protein and S*PARCL1* mRNA. **I** Cells within the VZ in (**H**) were clustered into four classes by FOXJ1 protein± and S*PARCL1* mRNA±. Cluster A: S*PARCL1*/FOXJ1-/-, B: S*PARCL1*/FOXJ1 +/-, C: S*PARCL1*/FOXJ1-/+, D: S*PARCL1*/ FOXJ1 +/+ (Fig. S5C, S5D, Table S4). See **Materials and Methods** for clustering. Scale bar = 20 μm.

Moreover, a fraction of tRG belongs to the state 7 (astrogenic state) in the pseudo-time trajectory (**Fig. 4E, 4F**). We examined whether these tRG cells differentiate into astrogenic cells. We chose *FOXJ1* as an ependymal marker and *SPARCL1* as an early astroglial marker based on a recent study in mice (Liu *et al*, 2022). While *SPARCL1* is expressed in all states generated by NPC2 (**Fig. S6B**), *SPARCL1* expression is highest in astrogenic tRG among the three tRG subgroups, whereas *FOXJ1* expression is nearly exclusive to ependymal tRG (**Fig. 4F**). Furthermore, cells expressing high levels of *SPARCL1* do not express *FOXJ1* (**Fig. 4G, Table S3**). These features of the astrogenic tRG are essentially the same as those of the astrogenic state (state 7) of the pseudo-time trajectory (**Fig. 4D**). Consistently, whereas FOXJ1^+^ cells were located close to the apical surface, those expressing *SPARCL1* were observed relatively far from the apical surface in the VZ (**Fig. 5H, 5I, S6C, Table S4**). Taken together, our analyses strongly suggest that tRG cells adopt both the ependymal and astroglial fates.

### Ferrets and humans show a homologous developmental transcriptomic profile of progenitor subtypes

Both ferrets and humans are positioned in distinct phylogenetic branches; both represent complex brain features of gyrencephalic mammals. Therefore, we compared our temporal series of ferret single-cell transcriptomes with a previously published human dataset (Nowakowski et al, 2017) to examine which processes are species-specific or common for the two species. We merged the two datasets by pairing mutual nearest neighbor cells (MNN) following canonical correlation analysis (CCA) across species (Stuart and Butler et al, 2019) and found that many cell types, including RG, IPC, OPC, and neurons, were clustered together (**Fig. 6A, S7A**). Temporal patterns and variety of neural progenitors during cortical developments were similar to each other between humans and ferrets at the single cell transcriptome level. Nonetheless, the timescales of cortical development significantly differed: on the UMAP plot, ferret E25 cells closely distributed with human GW8 cells; ferret E34, with human GW11–14; ferret E40–P1, with human GW15–16; and ferret P5–P10, with human GW17–22 (**Fig. 6B**).

**Figure 6.**
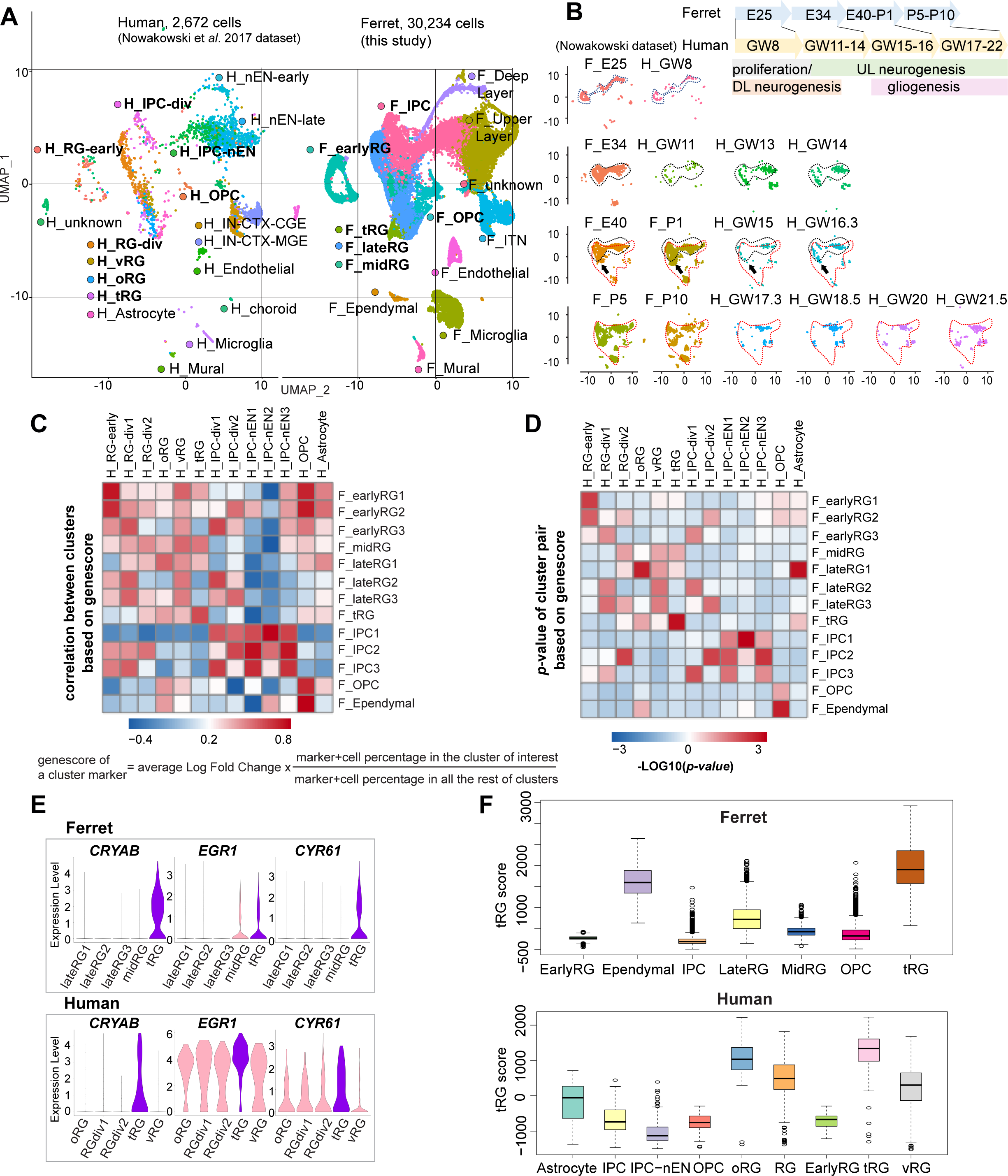
Comparison of molecular identity of RG subtypes between ferret and human. **A** UMAP visualization of integrated human (n = 2,672; left) and ferret (n=30,234; right) single-cell datasets colored according to the different clusters. The names of clusters from human and ferret cells begin with “H” and “F,” respectively. **B** Homologous temporal pattern of transcriptomic characters of progenitors and their progenies between ferrets and humans. Corresponding neurogenic stages between ferrets and humans can be assigned by cell distribution at homologous positions in UMAP plots. Gliogenic RG cells (a subtype of the “late_RG” group and OPC in **Fig. 2A**) were first distinguished transcriptionally at ferret E40 and at human GW14, as pointed out by arrows. **C, D** Correlation coefficient (**C**) and significance (**D**) between indicated clusters of ferrets and humans, calculated by marker gene scores. See **the text** and **Materials and Methods** for details. **E** Normalized expression levels of the indicated genes in humans and ferrets from each progenitor cluster. See **the text** and **Materials and Methods** for details. **F** tRG scores of the indicated clusters are presented as box and whisker plots for humans and ferrets. As for the definition of cluster scores, see **the text** and **Material and Methods**. These represent the range without outliers (lines) and upper and lower quartiles with the median (box). Outliers are represented as points outside the box. Outliers are 1.5-fold larger or smaller than interquartile range from the third or first quartile, respectively.

Next, we compared the similarities among individual RG subtypes across species. To quantify the features of an RG subtype, we introduced the parameter “genescore” (Bhaduri et al, 2020), a parameter that reflects the enrichment and specificity of a marker gene for a given cluster; the genescore for a particular marker gene is defined by multiplying the average enrichment of expression quantity in cells of the cluster (fold change) and the ratio of the number of marker-expressing cells in the cluster to the cell number in all other clusters (see **Materials and Methods**). Comparison of genescores of two arbitrary RG subtypes between two species revealed a high correlation in the early and late RG clusters (vRG, and oRG for the human dataset) across species (**Fig. 6C, 6D, Table S5**).

tRG cells also showed a remarkable similarity between the two species (**Fig. 6C, 6D**), as represented by a high level of expression for the combination of *CRYAB*, *EGR1,* and *CYR61* (**Fig. 6E**). To better examine this similarity, we defined the “cluster score” for individual cells as a linear combination of the expression level of marker genes in a cell, which is weighted by its genescore in the human cluster of interest. Calculation of the cluster score for ferret tRG using genescores of the human tRG cluster resulted in a higher score than any other cluster in both datasets (**Fig. 6F**). These results confirm, via transcriptomics, that the ferret tRG cluster is very close to the human one.

### Transcriptional analysis of human tRG subtypes by integration with ferret tRG subtypes

Next, we assessed whether human tRG cells also possessed ependymal and gliogenic potential, as suggested in ferret cells. We chose a recently published human dataset (Bhaduri et al, 2021) for comparison, because this study containing GW25 dataset which included more tRG cells than previous studies that did not contain GW25 data. Furthermore, we used only data at GW25. After excluding neurons and other cell types from the analysis, we identified respective human clusters for tRG and oRG cells based on their marker gene expression (**Fig. 7A, Table S6).** Then, we merged this human dataset and ferret NPCs (**Fig. 7B, 7C**) by the MNN after CCA as described before when we merged the entire series of progenitor samples of ferrets and humans (**Fig. 6A)**. This procedure revealed that human tRG and oRG share similar transcriptomes with ferret tRG and late RG that must include oRG, respectively (**Fig. 7D-H and Fig. S8A, S8B**). It is worth noting that early RG in ferrets showed the highest similarity with “OLIG1” in human data, and no cell type in human data corresponded to ferret “midRG,” likely because only GW25 cells were used for comparison (**Fig. S8A, S8B**).

**Figure 7.**
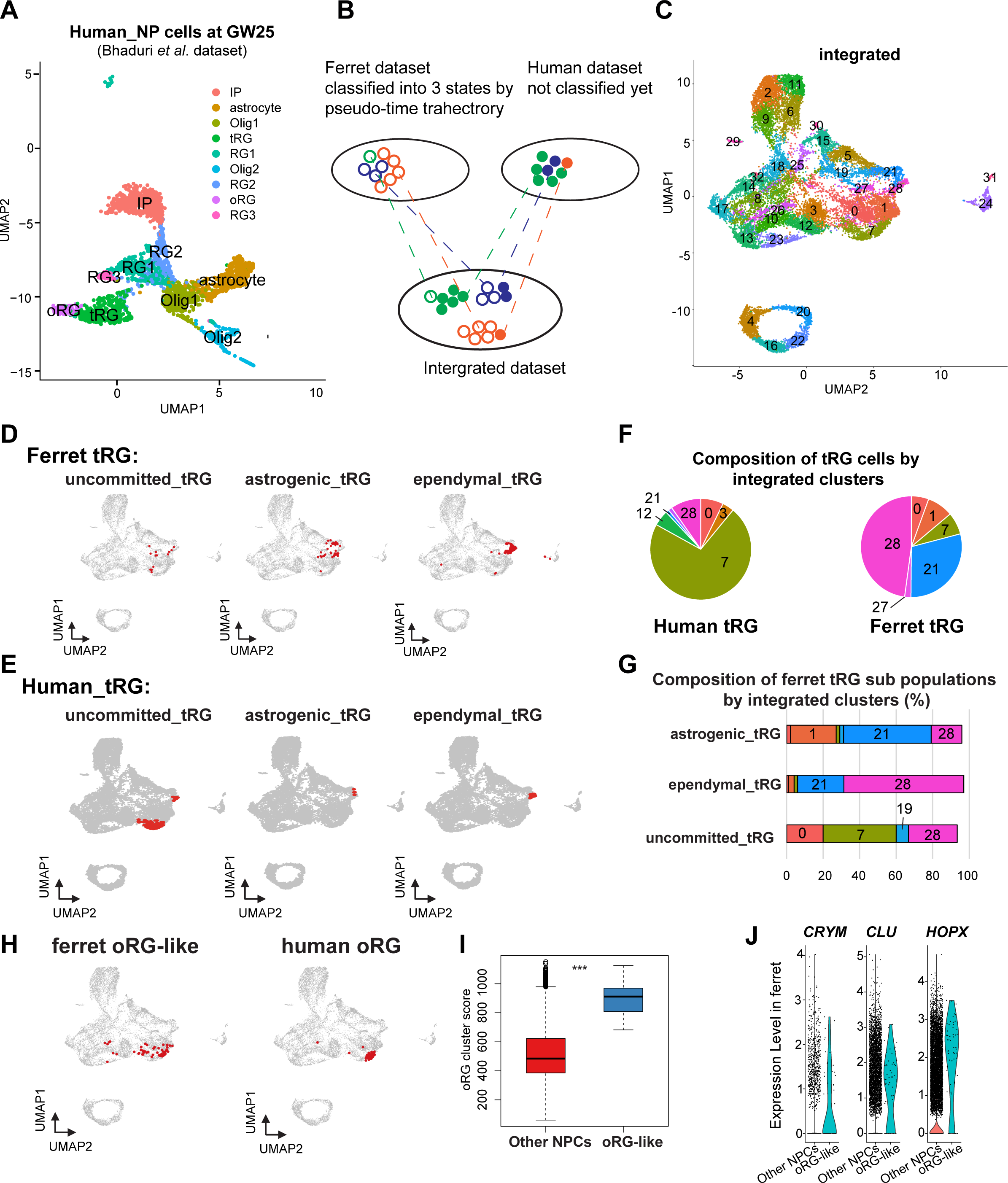
- Identification of tRG subtypes in humans and oRG in ferrets. **A**UMAP visualization of human brain cells at GW25 (and removing neurons and other cell types). Cells are colored by cell type and identified by marker genes (data from Bhaduri et al, 2021). **B** Schematic of the integration strategy of human and ferret subtypes. We merged the two datasets by pairing mutual nearest neighbor cells (MNN) following canonical correlation analysis (CCA) across species (Stuart and Butler et al, 2019). **C** UMAP visualization of integrated ferret and human datasets colored and numbered by different clusters. **D**, **E** Identification of three tRG subtypes in ferrets (D) and humans (E). The red dots highlight the indicated tRG subtypes named after pseudotime trajectory analysis in **Fig. 4**: presumptively “uncommitted”, “astrogenic” and “ependymal”. **F** Distribution of humans (left) and ferrets (right) tRG in the integrated dataset. Human and ferret tRG were identified from the separated dataset. The numbering corresponds to cluster numbers in the integrated clustering (C) of human and ferret datasets. **G** Distribution of ferret tRG subtypes (identified by the pseudo-time analysis) in the integrated dataset. A major cell population of each of the 3 ferret tRG subtypes (classified by the pseudo-time analysis), more or less, belongs to a single cluster in ferret-human integrated clustering; astrogenic tRG corresponds to cluster 21; ependymal tRG to cluster 20; uncommitted tRG to cluster 7. **H** Identification of oRG-like cells in ferret (left) and oRG cells in human (right), by the same way that we have done for tRG. All cells highlighted as red dots. **I** oRG scores for oRG-like cells and other NPCs in ferrets. The result indicates that oRG-like cells share more transcriptomic features than other NPCs, whereas ordinary clustering among ferret NPCs failed to distinguish oRG-like cells from the rest. Box and whisker plots indicate ranges (lines) and upper and lower quartiles with the median (box). **J** The expression levels of *CRYM*, *CLU*, and *HOPX* in oRG-like cells and other NPCs in ferrets.

The integration of ferret and human datasets showed that tRG from both species were mainly distributed into three different clusters in the UMAP space: 7, 21, and 28 (**Fig. 7F, S8C**). Cluster 7 cells highly expressed late onset RG genes (such as *APOE*, *FABP7*, *NOTCH2*, and *DBI*; **Fig. S8D**), suggesting that tRG cells in cluster 7 were presumably at a late RG-like state, being not yet committed to neither astrogenic nor ependymal fates (mostly in the NPC 3 state in the ferret pseudo-time trajectory; **Fig 3A, 3E**). In contrast, cells in clusters 21 and 28 highly expressed marker genes for the astrocyte (*GFAP* and *AQP4*), and the ependymal (*FOXJ*1) clusters, respectively (**Fig. S8D, Table S6**). Then, we confirmed that the three states of ferret tRG cells, committed to the astroglia, ependymal fates and uncommitted states, were largely assigned to clusters 21, 28 and 7, respectively (**Fig. 7G**), guaranteeing the reliability of this method. Thus, our results raise the possibility that tRG cells in humans possess the potential to generate astrocytes and ependymal cells as suggested for ferret tRG. This analysis also reveals that the proportion of the 3 tRG subtypes was very different between human and ferret datasets (**Fig. S7D–F**); the two presumptively committed tRG are very minor populations in humans, possibly due to the differences in developmental stages where tRG of these subtypes are enriched (see Discussion).

### Human cortical organoids lack tRG population

Human cortical organoids are a powerful model for understanding cortical development (Eiraku *et al*, 2008). It is, therefore, important to know how developing ferret cortices resemble human cortical organoids especially regarding to features shared by human tissues and ferrets. We then decided to analyze two different datasets of cortical organoids to see whether tRG is present in those organoids (**Fig. S9A-D** for Bhaduri et al., 2020; **Fig. S9E-F** for Herring et al., 2022), by focusing on CRYAB-expressing cells (**Fig. S9**). We assigned cells according to the original annotations, which included the information of organoid lines, age, and state (**Fig. S9A**). Our clustering resulted in 32 clusters where CRYAB-expressing cells were detected in two out of four cell lines (**Fig. S9B**). We noted that a tRG cluster was absent in the original annotation. To further confirm the absence of a tRG cluster in organoids, we integrated organoid dataset with the dataset derived from human primary tissues (Bhaduri et al., 2020). After the integration, we failed to find organoid-derived CRYAB-expressing cells that overlapped with tRG cells from the human primary tissues (Bhaduri et al., 2020; **Fig. S9C-D**), nor CRYAB-expressing cell itself in the other dataset (Herring et al., 2022; **Fig. S9E**). Our analyses thus indicate that tRG-like populations seem to be lacking in organoid datasets that are currently available (Bhaduri et al., 2020, Herring et al., 2022).

### Prediction of ferret oRG-like cells via the identification of cells homologous to human oRG cells

As mentioned before, oRG could not be identified as a separate cluster in the ferret dataset (**Fig. 1**). We attempted to assign oRG-like cells in the ferret dataset using human oRG cells as anchors via the integration of two datasets by the MNN after CCA (**Fig. 7B**). We could assign oRG-like cells, which were located near the human oRG cluster (**Fig. 7H**). To assess the degree of similarity to human oRG, we calculated the oRG score for each ferret oRG-like cell as we did for tRG (**Fig. 6F**). The assigned oRG-like cells had significantly higher oRG cluster scores than all other NPCs (**Fig. 7I**). Furthermore, we searched genes that were highly expressed in oRG-like cells by comparing oRG-like cells and other NPCs in the ferret dataset (**Table S6**). Consistent with the human dataset (Bhaduri *et al*, 2021), the expression of *HOPX, CLU,* and *CRYM* was higher in oRG-like cells than in other ferret NPCs (**Fig. 7J, S8E**). However, human oRG markers such as *HOPX* and *CLU* were also expressed by vRG and tRG in ferrets (**Fig. S3D, S8F**). Therefore, oRG at the transcriptome level can only be distinguished by the similarity with the whole transcriptome or a combination of several markers, but not by a few marker genes, whereas oRG can be unambiguously identified by its location in the tissue (**Fig. S1**).

## DISCUSSION

In this study, we provide a ferret dataset of single-cell transcriptomes of neural progenitors covering the entire cortical neurogenenesis and early gliogenic phase, revealing their diversity and temporal patterns during cortical development. Comparison of these features between ferrets and humans (Nowakowski *et al*, 2017; Bhaduri *et al*, 2021) indicated at a high resolution that these two gyrencephalic mammals shared a large proportion of progenitor variations and their temporal sequences despite their extremely different neurogenesis timescales. It is remarkable that tRG is conserved between these two phylogenetically distant species. A ferret model allows us to identify the mode of tRG generation, to predict its descendant fates and to analyze cell dynamics in tissues. These findings have not been realized by previous studies, re-evaluating ferrets as a valuable model species to study the neural development of gyrencephalic animals. This comparison between humans and ferrets could not be performed unless genomic information, especially genome-wide gene models, was greatly improved; there had been sufficient genomic information accumulated for ferrets, but mainly for coding regions for genes, which had not been well connected to their 3’-UTR. However, to carry out a large scale of single cell RNA seq quantitatively with minimizing the loss of transcript species, accurate 3’- UTR sequences and their link to the coding part are indispensable, because typical scRNA-seq platforms are based on oligo-(dT)-primed cDNA synthesis. We made tremendous efforts to get improved gene models that include 3’-UTR sequences, making it possible for us to compare single cell transcriptome of ferret cortical progenitors with a huge volume of single cell transcriptome datasets with high qualities, including those for humans and mice.

### The fate of tRG progenies

Our pseudotime trajectory analyses and immunohistochemistry analyses suggested that ferret tRG cells differentiate into ependymal cells and astrogenic cells. Although the number tRG cells used in the trajectory analysis was limited (167 cells; **Fig. 3**), three tRG subtypes, which are presumably “precommitted”, “ependymal” and “astrogenic” tRG also formed three distinct territories within the original tRG cluster that consisted of 409 cells on UMAP, indicating a consistent heterogeneity within tRG cluster. This mode of the presumptive ependymal cell formation is likely parallel to that of the bifurcation of ependymal cells and adult neural stem cells from the same progenitor population in the striatum on the ventral side (Ortiz-Álvarez *et al*, 2019).

### Comparison of human with ferrets in corticogenesis

Our cross-species comparative analysis predicts that tRG in humans also differentiate into ependymal and astroglial cells although there is no *in vivo* evidence yet (**Fig. 6E**). Furthermore, the rare presence of two presumptively committed tRG subgroups in humans makes this prediction tentative at present. If this prediction is the case, the uncommitted tRG fraction is dominant in GW25 human brains unlike in ferret brains, suggesting that human tRG at GW25 progresses along the differentiation axis less than ferret tRG at P5–P10, most of which are already committed to either ependymal or astrogenic states.

While GW25 has been almost at the latest stages experimentally available from human embryonic brain tissues due to ethical reasons, Herring et al. recently reported a large dataset that spans late embryonic to postnatal development in human prefrontal cortex, in which we were not able to detect a tRG population nor other progenitor subtypes, as well as an ependymal cell type (Herring *et al*, 2022), possibly due to regional differences in the collected samples from our ferret dataset. Thus, this postnatal dataset does not seem to be appropriate to compare with our ferret dataset or even human prenatal dataset (Bhaduri et al. 2020) to follow tRG progenies (**Fig. S9**). More enrichment of datasets in the human lateral cortices at the developmental stages after GW25 onward are necessary to test whether our results obtained from ferrets regarding tRG progeny fates can be generalized.

### Commonalities and differences among characteristics of neural progenitors between human ferrets

In this study, our transcriptomic data supported the notion that ferrets are a good model system to investigate mechanisms commonly (at least in both primates and carnivores) underlying the cortical development in gyrencephalic mammals. At the same time, we acknowledge the differences in corticogenesis between ferrets and humans. Most prominently, the time scale of corticogenesis progression is greatly different between the two species, while the variety of NPCs and their temporal patterns of appearances and differentiation exhibit similarities. In addition, the organization of cortical scaffold seems differ from each other; in human corticogenesis, two major germinal layers, the VZ and OSVZ, become segregated during the expansion phase of RG in the neurogenic stage, due to the loss of vRG, which extend from the ventricular surface to the laminar surface, passing through the OSVZ (Nowakowski *et al*, 2016). In contrast, ferrets maintain vRG with a long radial fiber in the VZ alongside with tRG cells, even beyond the expansion phase of tRG (**Figure 3A, S4G**).

### Models as human cortical development

Our preliminary analysis of two datasets from human cortical organoids suggests the lack of tRG-like cells and potentially their progeny cell types (**Fig. S9**, Bhaduri et al., 2020; Herring et al., 2022). This situation might be improved by optimizing culture protocols. If tRG and ependymal cells, one of the presumptive tRG progeny, are generated in human organoids, as we found in ferrets, it would represent a significant advancement in the human organoid model because ependyma are a fundamental structure for ventricular formation. Further studies using human cortical organoids are expected to be able to generate tRG, and to examine its progeny fates.

In ferrets, genetic manipulations can be achieved through *in utero* or postnatal electroporation, as well as via virus-mediated transfer of DNA (Borrell, 2010; Kawasaki *et al*, 2012; Matsui *et al*, 2013; Tsunekawa *et al*, 2016). Thus, it is theoretically possible to disrupt the *CRYAB* gene *in vivo* in ferrets to investigate its role in tRG and their progeny, including ependymal cells, and to track the tRG lineage. If the *CRYAB* gene is essential to form ependymal layers, we will be able to explore how the ventricle contributes to cortical folding and expansion. Despite our extensive efforts over a year, we have thus far been unsuccessful in knocking in and/or knocking out the *CRYAB* gene. Nevertheless, we anticipate that technical advances will surpass our expectations, both in ferret and human organoids. Taken together, these functional studies in ferrets as well as in human organoids hold promising insights into the understanding of the tRG lineage and its contribution to cortical development in the near future.

## MATERIALS AND METHODS

## EXPERIMENTAL MODEL AND SUBJECT DETAILS

### Ferret brain tissue samples

Pregnant ferrets were purchased from Marshall BioResources. We also bred them to maintain a population that is sufficient to regularly get pregnant females in the animal facility of RIKEN Center for Biosystems Dynamics Research under the license given from Marshall BioResources. All procedures during animal experiments were performed in accordance with the legal and institutional ethical regulations of RIKEN Center for Biosystems Dynamics Research.

Ferret pups are born at day 41 or day 42 of gestation. The day of birth was counted as postnatal day 0 (P0). Ferret embryos and pups were euthanized prior to brain removal.

Samples used for single cell RNA-sequencing were obtained from 1 or 2 pups for each developmental stage: embryonic days 25, 34 and 40 and postnatal days 1, 5 and 10. Samples used for immunohistology-based quantifications were obtained from 2 or 3 pups from different mothers. Samples used for a time-lapse imaging were obtained from 1 animal per experiment, which is performed 4 times, independently.

## METHOD DETAILS

### Gene model construction of the ferret genome

In this study, we used a ferret genome reference with expanded reference gene annotations with DDBJ (detail see below). Briefly, gene models were constructed using Chromium by tagging all fragments from a long genomic DNA in a droplet, so that sequences from long genomic DNAs could be successfully aligned to cover so far unconnected genomic regions in the NCBI (NCBI Assembly ID: 286418 (MusPutFur1.0)) database. Detailed information after mapping is listed in the **Supplementary Table S1B**.

### RNA-seq library preparation and sequencing

Total RNA was extracted from the embryonic tissues (Supplementary Table S7) using the RNeasy mini kit (Qiagen). Concentration and length distribution of the RNA were checked with the Qubit RNA HS Assay Kit on a Qubit 2.0 Fluorometer (Thermo Fisher Scientific) and the RNA 6000 Nano Kit on a 2100 Bioanalyzer (Agilent Technologies). Sequencing libraries were prepared using 1 μg of total RNA with the TruSeq Stranded mRNA Sample Prep Kit (Illumina) and the TruSeq single-index adaptor (Illumina), except for the sample ‘gonad and mesonephros’ (see Supplementary Table S7) for which 500 ng of total RNA was used. The RNA was fragmented at 94°C for either 8 minutes or 2 minutes, which resulted in libraries with variable insert size distributions (Hara *et al*, 2015). The optimal numbers of PCR cycles were pre-determined using the KAPA Real-Time Library Amplification Kit (Roche) as described previously (Tanegashima *et al*, 2018). Libraries were sequenced with the Rapid Run mode of HiSeq1500 (Illumina) using the HiSeq PE Rapid Cluster Kit ver. 2 (Illumina) and the HiSeq Rapid SBS Kit v2-HS (Illumina) to produce paired-end reads of 127 nucleotides. Base calling was performed with RTA ver. 1.18.64, and the fastq files were generated with bcl2fastq ver. 1.8.4 (Illumina).

### Genomic DNA extraction and Chromium genome library construction

Genomic DNA was extracted from a liver of an adult female ferret using the CHEF Mammalian Genomic DNA Plug Kit (BioRad, Cat. No. #1703591). In brief, the liver tissue of about 50 mg was homogenized in PBS (-) with a Dounce tissue grinder on ice, fixed with ethanol, and subsequently embedded in low melting point agarose gel (BioRad). After the protein digestion with Proteinase K (QIAGEN, Cat. No. #158920) and RNA digestion with RNase A (QIAGEN, Cat. No. #158924), the gel plugs were further digested by Agarase (Life Technologies, Cat. No. #EO0461). DNA was purified by the drop dialysis method using the MF-Millipore Membrane Filter (Merck Millipore, Cat. No. #VCWP04700). Length distribution of the genomic DNA was analyzed by pulsed-field gel electrophoresis, which exhibited an average length of over 2 Mbp. The Chromium library was constructed using the Chromium Genome Reagent Kit v1 Chemistry (10x Genomics, Cat No. #PN-120216), and sequencing was performed with an Illumina HiSeq X to obtain paired-end 151 nt-long reads.

### Genome assembly

De novo genome assembly using the Chromium linked-reads was performed with Supernova ver. 2.0.0 (10x Genomics) with the default parameters, and the resultant sequences were output with the option ‘pseudohap’. Detection and masking of repetitive sequences were performed using RepeatMasker ver. 4.0.7 (Smit et al, 2013-2015) with NCBI RMBlast ver. 2.6.0+ and the species-specific repeat library from RepBase ver. 23.01.

### Gene model construction

Gene models for gene expression level quantification were constructed in the three following steps. First, ab initio gene prediction was performed by BRAKER ver. 2.0.5 (Hoff *et al*, 2019). In this process, GeneMark-ET ver. 4.33 (Lomsadze *et al*, 2014) with the information of spliced RNA-seq alignment was used for training, and AUGUSTUS ver. 3.3 (Stanke *et al*, 2008) with the parameters of ‘UTR=on, species=human’ was used for gene prediction. For the abovementioned training of BRAKER, the RNA-seq reads were aligned to the repeat-masked genome sequences by STAR ver. 2.5.4a (Dobin *et al*, 2013) with default parameters using the entire set of RNA-seq reads of various tissue types (Supplementary Table S7). Second, the information of RNA-seq alignment was directly incorporated into the gene models to improve the coverage of the 3’ UTR region of each gene which is not reliably predicted in ab initio gene prediction. This computation was performed with HISAT2 ver. 2.1.0 (Kim *et al*, 2019) and StringTie ver. 1.3.4d (Kovaka *et al*, 2019), whose output GTF file was merged with the gene models produced by BRAKER using the merge function of StringTie. Finally, the existing transcript sequences of MusPutFur1.0 available at NCBI RefSeq (GCF_000215625.1) was incorporated into the gene models by mapping their sequences to the repeat-masked genome sequences using GMAP ver. 2017-11-15 (Wu & Watanabe, 2005) with the ‘not report chimeric alignments’ option. The gene name of each locus was adopted from the annotation in RefSeq, and the gene name of a newly predicted locus was assigned according to the results of a BLASTX search against the UniProtKB Swiss-Prot database release 2020_01. The assignment was performed only for genes with a bit score (in the abovementioned BLASTX search) of greater than 60. When a locus has multiple transcripts, the one with the highest score was adopted for gene naming. If a transcript estimated by ab initio prediction bridged multiple genes, they were incorporated into the gene models as separate genes.

### Completeness assessment of assemblies

To assess the continuity of the genome assembly and gene space completeness of the gene models, gVolante ver. 1.2.1 (Nishimura *et al*, 2017) was used with the CEGMA ortholog search pipeline and the reference orthologues gene set CVG (Hara *et al*, 2015).

### *In utero* and postnatal electroporation

*In utero* electroporation (IUE) and postnatal electroporation in ferrets was performed as described previously (Kawasaki *et al*, 2012; Matsui *et al*, 2013; Tsunekawa *et al*, 2016 for IUE; Borrell, 2010 for postnatal electroporation) with modifications. Briefly, pregnant ferrets or ferret pups were anesthetized with isoflurane at indicated stages of the development. The location of lateral ventricles was visualized with transmitted light delivered through an optical fiber cable. Three µl of plasmid DNA solution was injected into the lateral ventricle at indicated developmental stages using an injector. Each embryo or pup was placed between the paddles of electrodes, and was applied a voltage pulse of 45V at the duration from 100 msec to 900 msec (100,0 Pon and 1000,0 Poff) 10 times for in utero electroporation, and under the same condition except a voltage of 60V for postnatal electroporation (CUY21 electroporator, Nepa Gene).

### Plasmids

To assure a stabilized expression of transgenes, we combined expression vectors with a hyperactive piggyBac transposase system to integrate expression vectors into the genome for a stable expression (Yusa *et al*, 2011). To enrich neural progenitor cell populations in “AG” samples used for the single-cell transcriptome analysis, pLR5-Hes5-d2-AzamiGreen (0.5 µg/µl; Hes5 promoter was gifted from Kageyama Laboratory; Ohtsuka *et al*, 2006) was electroporated at E30 for E40, or at E34 for postnatal samples, together with pCAX-hyPBase (0.5 µg/µl). Among them, for P1 and P10 samples, pPB-LR5-mCherry (0.5 µg/µl) was also included in the DNA solution.

To sparsely label the cell cytoplasm for detailed imaging of individual cells on vibratome-cut thick cortical sections, we have used pPB-LR5-floxstop-EGFP combined with a low concentration of Cre-expressing plasmid. The concentrations used for these plasmids were 0.5 µg/µl or 1.0 µg/µl for a GFP labeling and 1 ng/µl or 10 ng/µl of Cre-expressing plasmid for P0 or P5 samples, respectively (Figure 2A). For all electroporation experiments, Fast Green solution (0.1 mg/ml; Wako Pure Chemical Industries) was added into the freshly prepared mixture of plasmid DNA to visualize the injection.

For in situ hybridization probes, the PCR product was inserted into pCR-BluntII-TOPO for cloning and sequencing.

### Library preparation for scRNA-seq

10x v2 Chromium was performed on dissociated single cells according to the manufacturer’s protocol.

### Single cell isolation and sorting of Azamigreen-expressing cells

Cell suspension was prepared as reported previously (Wu *et al*, 2022) with modifications. Brains were collected at indicated stages, transferred in the ice-cold dissection Dulbecco’s Modified Eagle Medium (DMEM) F12 and meninges were removed. Somatosensory area was dissected from embryonic or postnatal cortices using an ophthalmic knife (15°) under a dissection microscope. Sliced tissues were dissociated via enzymatic digestion with papain at 37℃ for 30 - 45 min in ice-cold Hank’s Balanced Salt Solution (HBSS) (-) with EDTA (0.1 M). Dissociated cells were centrifuged at 1000xg for 5 min to remove papain by washing with PBS and were resuspended in 0.375% BSA/HBSS (-), or in the sorting buffer (DMEM F12 + GlutamaX (Thermo Fisher); 0.1 % of Penicillin/Streptomycin (Millipore); 20 ng/ml human basic FGF, Peprotech; 1xB27 RA-, Gibco) for cell sorting (see below). Homogenous cell suspension filtered through 35 µm strainer (Falcon).

To concentrate neural progenitor cell (NPC) populations at the expense of mature neurons, two methods were used for each sample, except for E25, from which cells were collected using whole somatosensory area:

1. “T” samples: the cortical plate (CP) of cortical sections were dissected out and a part of the intermediate zone (IZ) was likely to be included in the discarded region.
2. “AG” samples: brains collected at indicated stages were electroporated in utero by azamiGreen (AG)-expression vector under the control of Hes5 promoter. The latter method allows the expression of AG in only NPC by the presence of a degradation signal “d2” in the vector to degrade AG protein in HES5-negative differentiating progeny. Only “AG” samples were processed with cell sorting and dissociated cell obtained as described above were placed in a SH800 cell sorter (SONY) to sort azamigreen-positive cells in 0.375%BSA/HBSS(-) solution. Cell survival and cell number were quantified by Countess or Countess II (Invitrogen), prior to an application of single cell isolation using 10x v2 Chromium kit.

Samples from the same developmental stage, except for P10 “T” sample, were born from the same mother and were collected by applying either of above methods on the same day and processed in parallel.

### Library preparation and Sequencing

Single-cell libraries were generated according to the manufacturer’s instructions. Cell suspensions were diluted for an appropriate concentration to obtain 3,000 cells per channel of a 10X microfluidic chip device and were then loaded on the 10X Chromium chips accordingly to the manufacturer’s instructions.

Total cDNA integrity and quality were assessed with Agilent 2100 Bioanalyzer.

Libraries were sequenced on the HiSeq PE Rapid Cluster Kit v2 (Illumina), or the TruSeq PE Cluster Kit v3-cBot-HS, to obtain paired-end 26 nt (Read 1)-98 nt (Read 2) reads.

### Immunohistology and confocal imaging

Ferret brains were removed from embryos or pups and fixed for 1 or 2 overnights, respectively, in 1% paraformaldehyde (PFA) prepared in 0.1M phosphate buffer (PB, pH 7.4) at 4℃. P35 ferrets were transcardially perfused with cold PBS, followed by 4% PFA, under deep anesthesia with isoflurane, then collected brains were post-fixed with 1% PFA. After the fixation of brains or cortical tissue slices after a live imaging, they were equilibrated in 25% sucrose overnight before embedding in O.C.T. compound (TissueTek, Sakura) and frozen in liquid nitrogen. Frozen samples were stored at -80℃ prior to a coronal sectioning using a cryostat (CM3050S Leica Microsystems, 12µm sections). After equilibration at room temperature, sections were washed in PBS with 0.1% Tween (PBST), followed by a treatment with an antigen retrieval solution of HistoVT one (Nacalai Tesque) diluted 10 times in milliQ water, at 70℃ for 20 min. Sections were then blocked 1 hr at the room temperature (RT) in PBS with 2% Triton-X100 and 2% normal donkey serum (Sigma), followed by an incubation overnight at 4℃ with primary antibody diluted in the blocking solution. After washing in PBST three times, sections were treated with appropriate fluorescence-conjugated secondary antibodies (1:500) along with 4’,6-diamidino-2-phenylindole (DAPI, 1:1,000) for 1 hr at room temperature. Sections were washed again prior to mounting with PermaFluor solution (Therma Fisher Scientific).

Immunostaining of thick ferret brain sections (200 µm) were performed as previously reported (Tsunekawa et al., 2016). Briefly, ferret brains were fixed in 1%PFA, washed in PB overnight at 4℃, and embedded in 4% low-melting agarose (UltraPure LMP agarose, Thermo Fischer Scientific). Embedded brains were sliced coronally at 200 µm thickness by a vibratome (LinerSlicer, DOSAKA EM) on ice. Floating sections were washed three times with PBST, treated with the blocking solution for 1 hour at RT and incubated with primary antibodies for 5 or 6 overnights at 4℃ under shaking. Sections were then washed in PBST and treated with secondary antibodies for 5 or 6 overnights at 4℃ under shaking. After washing, we mounted brain slices with CUBIC solution 2 to allow the transparency.

Fluorescent images were acquired using an FV1000 confocal microscope (Olympus, Japan). Somatosensory cortices were captured with 20X or 100X objective lenses. When 100x objective lens was used, z-stacks of confocal images were taken with an optical slice thickness 1.0 µm or 1.5 µm and MAX-projection images obtained by FiJi are shown.

All the antibodies used in this study are listed in Key Resources Table, and are used at following concentrations: chicken anti-EGFP, 1:500; mouse anti-Alpha B Crystallin, 1:500; Rabbit anti-FoxJ1, 1:500; goat anti-Olig2, 1:500; rat anti-Eomes, 1:500; rabbit anti-Pax6, 1:500; mouse anti-Pax6, 1:500; mouse anti Gfap, 1:500; rabbit anti-HopX, 1:500; rat anti-RFP, 1:500; goat anti-adenine cyclase III, 1:250; rat anti-Ctip2, 1:500; mouse anti-Satb2, 1:500, rat anti-Ki67, 1:500.

### Probe preparation

Primers for PCR targeting ferret *CLU* gene were designed with Primer3 ver 0.4.0 software. Total RNA was isolated from embryonic ferret brain, collected in Trizol. cDNA was generated from total RNA using the Prime Script 1st strand cDNA synthesis kit (Takara) according to the manufacturer’s recommended procedure. Target cDNA was inserted into pCR-TOPOII-Blunt plasmid for cloning and sequencing using M13F or M13R primers. Anti-sense cRNA probes were then generated by in vitro transcription using T7 promoter.

### *In situ* hybridization

ISH was performed as described previously, with some modifications (Mashiko *et al*, 2012). Briefly, ferret brains were perfused in 4%PFA in PBS and were fixed at 4%PFA in 0.1M PB, Fixed brain were incubated in 30% sucrose/4%PFA in 0.1M PB for at least 1 day and were stored at -80℃ until ISH. Frozen brains were sectioned in coronal plane at 25 µm using cryostat (CM3050S Leica Microsystems). Sections on slides were post-fixed at 4% PFA in 0.1M PB and treated with proteinase K (Roche). Sections were hybridized with digoxigenin (DIG)- labeled probes at 72°C overnight in hybridization solution. Sections were then washed and blocked with donkey serum and incubated with pre-absorbed DIG antibody, conjugated to Alkaline phosphatase for 2 hours at room temperature. Color development was performed in solution containing NBT/BCIP (Roche).

## QUANTIFICATION AND STATISTICAL ANALYSIS FOR scRNA-seq

Statistical details including experimental n, statistical tests and significance are reported in Figure Legends.

### Alignment and raw processing of data

Fastq files were obtained from individual samples and were processed using Cell Ranger pipeline v2. Alignment was done using “Cell Ranger count” function with default parameters accordingly to manufacturer’s instructions to map reads to the ferret reference (MPF_Kobe 2.0.27). The raw data for each set of cells within a sample was obtained by cellranger count function and was read using Seurat “Read10X” function (Seurat v3.1.5) (Stuart *et al*, 2019), creating a matrix for unique molecular identified (UMI) counts of each gene within each cell.

### Filtering and normalization

Filtering and normalization were performed using Seurat (v3.1.5) (Stuart *et al*, 2019). Briefly, each sample was filtered by removing low-quality cells with unique feature counts less than 200, and genes expressed in less than 3 cells (Seurat function “CreateSeuratObject”). Different samples were then merged using the Seurat “merge” function. Merged data was further subset by keeping cells with features (genes) over 200 and less than 5000. Raw UMI counts were then normalized using “LogNormalize” as normalization method, which divided reads by the total number of UMIs per cell, then multiplied by 10,000 (Seurat “NormalizeData” function). This resulted in a total of 30,234 cells and 19,492 genes: E25 (3486 cells), E34AG (3223 cells), E34T (2260 cells), E40AG (1102), E40T (1581 cells), P1AG (2681), P1T (3641), P5AG (2429), P5T (2926), P10AG (3010) and P10T (3895).

### Single-cell clustering and visualization

Cell clustering was employed to the entire cell population after removing low-quality cells, using Seurat (v3.1.5) (Stuart *et al*, 2019). 2,000 highly variable genes were identified and used for the downstream analysis using Seurat “FindVariableFeatures” function (selection.method = “vst”), which allowed the calculation of average expression and dispersion for each gene. Normalized data was then processed for scaling using Seurat function “Scale Data” with default settings, which also allowed the regression of the batch. Principal component analysis (PCA) was performed with 2,000 variable genes to reduce dimensionality of the scaled dataset and 50 PCs were retained (Seurat “RunPCA” function). Clustering was then performed using graph-based clustering approach (“Find Neighbors” function) using top 20 PCs, which were selected based on the standard deviation of PCs on the elbowplot created by Seurat function “ElbowPlot” and based on statistical significance calculated by JackStraw application. Briefly, cells are embedded in a k-nearest neighbor (KNN) graph based on the Euclidean distance in a PCA space. Then, this knn graph is used to group cells on a shared nearest neighbor graph based on calculations of overlap between cells with similar gene expression patterns (Jaccard similarity). Cells were then clustered by Louvain algorithm implemented in Seurat “ClusterCells” function (resolution = 0.8, dims = 1:20). Next, we used Seurat “RunUMAP” function, which resulted in cell clusters being separated in embedding space while preserving the balance between local and global structure.

### Cluster annotations

Cluster markers were obtained using “FindAllMarkers” Seurat function. We tested genes that showed at least a 0.25-fold difference between the cells in the cluster and all remaining cells, and that were detected in more than 25% of the cells in the cluster. Clusters, which we called subtypes, were then annotated by comparing cluster markers to previously identified cell type markers in the literature for mouse and human datasets. The full list of markers is given under Table S1. The heatmap in **Fig. 2B** shows the expression data for the top 10 highly expressed marker genes for each cluster using a downsampling of maximum 500 cells from each cluster. Plotting cells onto the UMAP plot by their batch indicated that batches did not influence clustering (**Fig. S2A**) in accordance with the differential expression of cluster markers (**Fig. 2B, S3A)**. When a cluster was enriched based on the developmental stage, the information was included in the annotation. For example, “early RG” cluster was enriched in E25 samples, and expressed both common radial glial cell markers shared with other RG clusters in our dataset, but also expressed early onset-genes reported in mouse studies (Okamoto *et al*, 2016; Telley *et al*, 2019), such as *HMGA2*, *FLRT3*, *LRRN1* (Table S1). Therefore, we named this cluster as “early RG” based on its age-dependent properties.

To achieve more precise clustering, cycling cells were identified with known markers implemented in Seurat package (S genes and G2M genes). Clusters enriched in S genes or G2M genes were identified, including early_RG2 (S), early_RG3 (G2M), late_RG2 (G2M), late_RG3 (S), IPC2 (S) and IPC3 (G2M) (**Fig. S3C**). Subtypes of RG and IPC clusters were further combined to facilitate the data representation (**Fig. 2B**).

### Differential gene expression analysis

Differential gene expression (DEG) analysis was assessed using “FindMarkers” Seurat function with default parameters.

### Pseudotime analysis

Monocle 2 was used to construct developmental trajectories based on pseudotime ordering of single cells (Qiu *et al*, 2017; Trapnell *et al*, 2014). We generated a subset of clusters using “SubsetData” Seurat function using the raw counts. Combined progenitor clusters (F_earlyRG, F_midRG, F_lateRG, F_tRG, F_IPC, F_OPC) were selected. All clusters known to be generated from a non-cortical origin including microglia, endothelial and mural cells, were removed prior the pseudotime analysis. The expression matrix and a metadata file that contained above cluster information defined by Seurat were used as input for monocle package (**Table S2**).

The “differentialGeneTest” function was used to select genes for dimensionality reduction (fullModelFormulaStr = ‘∼Subtype.combined’). Top 1,000 genes were then applied for cell ordering. The visualization of the minimum spanning tree on cells was obtained by Monocle2 “plot_complex_cell_trajectory” function. To visualize the ordered cells on Seurat’s UMAP plot, we extracted the branch information for cells and added as metadata of the merged Seurat object (“AddMetaData” Seurat function).

### Processing External scRNA-seq Datasets for comparisons and preprocessing

scRNA-seq data of developing human brain from Nowakowski *et al*. (2017), from Bhaduri *et al*. (2021) were used for cross-species comparisons (**Figures 6, 7, S7-S9**). All cells included in the analysis and cluster assignments from the provided matrix were mapped to the cell type assignments provided in the metadata of Nowakowski et al. dataset and the cell type labels were used as provided by the authors without modification. For Bhaduri et al. dataset, GW25 sample was used for clustering by Seurat package as described above. For both, we removed low quality cells and clusters annotated based on anatomical regions other than somatosensory cortex, and removed cells with less than 200 features. For Bhaduri et al. dataset, we filtered to cells with at least 200 features, less than 6,500 features and less than 5% mitochondrial genes. This resulted in 25,485 cells from around 180,000 cells in Bhaduri et al. dataset, and in 2,673 cells in Nowakowski et al. dataset.

For organoid datasets, Herring et al. (2022) and Bhaduri et al., (2021) were used (**Figs. S8, S9**) with default settings in Seurat. The data driven from organoids at 5-months, 9-months, 12- months and organoids at 3-weeks, 5-weeks, 8-weeks, 10-weeks were used for Herring and Bhaduri datasets, respectively.

### Integrated analysis of human and ferret scRNA-seq datasets

Human datasets published by Bhaduri *et al*. (2021) and Nowakowski *et al*. (2017) were used for integration analysis with our ferret dataset using Seurat canonical correlation analysis (Stuart *et al*, 2019). First, a Seurat object was created for all datasets as described previously. Briefly, each object was individually processed with a normalization and variable features were identified. For human datasets, we excluded cells or samples obtained from anatomical regions other than somatosensory cortex. Then, the integration anchors between human and ferret Seurat objects were identified using “FindIntegrationAnchors” Seurat function with default settings. The integration was then performed with “IntegrateData” Seurat function. Scaling, PCA and UMAP dimensional reduction were performed to visualize the integration results.

### Cluster correlation analysis between ferret and human clusters

Correlation analysis was performed as described by Bhaduri *et al*. (2020), with modifications. Briefly, lists of cluster marker genes obtained by “FindAllMarkers” Seurat function on individual datasets were extracted prior to an integration (Table S5A), except for Nowakowski et al dataset, for which the marker gene list was provided. A score based on the specificity and the enrichment of cluster marker genes was generated (defined as “Genescore”). A genescore value was obtained by the multiplication of the average fold change that represents the gene enrichment, with the percentage of the cells expressing the marker in the cluster (pct.1) divided the percentage of the cells from other clusters expressing the marker (pct.2: specificity). This function was employed on the marker genes for a human subtype of interest in the space of marker genes for ferret NPC (RG, IPC, OPC and EP) clusters. Genescores for both ferret and human were then obtained for all markers shared between human subtype of interest (i.e. oRG) and ferret NPC subtypes. We then applied cor.test function using Pearson method to estimate a correlation and the significance of the correlation for the obtained genescores between ferret and human samples. These resulting values were represented for NPC subtypes on heatmap plots.

### Prediction of oRG-like/tRG-like cells by cross-dataset analysis

The cell type marker genes for oRG and tRG cell types were extracted from each human scRNAseq datasets. To predict human oRG-like and tRG-like cells in ferret sample more precisely, we removed mature neuron, microglia, and endothelia clusters, as well as clusters assigned as unknown, from individual datasets for an integration between ferret and human NPC clusters using Seurat package as described in the above section (Stuart *et al*, 2019). After identification of anchors between human and ferret Seurat objects using “FindIntegrationAnchors”, we extracted the information of each pair of anchors, which including one ferret and one human cell. If the human cell in the pair belongs to oRG (tRG) cluster in human dataset, we consider the other ferret cell in the same pair as oRG (tRG)-like cell.

### Cluster score calculation

Based on a genescore calculated for marker genes as previously described, a cell-type predictive model was generated. For both datasets, our custom-made R scripts were applied to generate matrices using either human oRG markers or tRG markers using our integrated subsets. In the space of matrices reduced with corresponding human marker genes, a score we defined as “oRG score” or “tRG score” was created by multiplying human genescore for each marker genes with the expression values of these genes in all cells present in the integrated subset data. The results were represented by beeswarm and boxplot.

### Evaluation of immunolabeled cell number and statistical analysis

The counting frames on images of immunolabeled sections consisted of 150 µm of width regions of interests (ROIs). The numbers of immunolabeled cells within selected ROIs were manually counted using the “cell counter” tool of FiJi software (Schindelin *et al*, 2012). The VZ was identified by its cell density visualized by DAPI, and according to its thickness measured by a vertical length from the ventricular surface at indicated stages (100 µm at P5, 70 µm at P10). The proportions of immunolabeled cells positive for various markers were calculated using the summed data counted within all ROIs. The data obtained from lateral cortices of somatosensory area was averaged from 2 to 3 sections.

For CRYAB-positive cells, we only counted cells with nuclei which were visualized by DAPI staining. The numbers of positive cells were counted using two or three different animals.

Quantification was followed by Wilcoxon test to assess statistical significance and data on each section were considered as n = 1.

### Quantification of RNAscope in situ hybridization and immunostaining image

Images of RNAscope in situ hybridization and FOXJ1 immunostaining were captured by using an IX83 inverted microscope (Olympus) equipped with Dragonfly 200 confocal unit (Andor), Zyla Z4.2 sCMOS camera (Andor), and UPLXAPO 60XO objective lens (Olympus). Imaging was performed to satisfy Nyquist sampling (xy = 48 nm/pixel by using 2x zoom optic and z = 136 nm interval) and 3D deconvolution was performed by ClearView-GPU™ option of Fusion software (Andor). Default deconvolution parameter is used except for number of irritation (n = 24) and mounting media reflection index (RI = 1.46). For quantification of images, CellProfiler version 4.2.4 (Stirling *et al*, 2021) was used. To detect mRNA dots of RNAscope in situ hybridization images, a single Z section of deconvolved images were processed by EnhancingSpeckles module (10 pixels size) and then mRNA dots were detected by IdentifyPrimaryObjects module, adaptive Otsu two class method, using default parameter except for size of adaptive window (size = 10). Dot’s diameter less than 4 pixels were removed as a background noise. Cytoplasmic ROIs were generated from smoothed RNAscope in situ hybridization images by MedianFilter module (window size = 10) and manually drawn seed ROIs of DAPI staining images by using IdentifySecondaryObjects module (propagation adaptive Otsu two class method, using default parameter). FoxJ1 immunostaining raw images were measured by MeasureImageIntensity module. Quantified data were visualized and clustered by using RStudio (version 2022.07.1+554) with R (version 4.0.3). Each of mRNA count data and protein staining intensity data was log transformed and normalized, and hierarchical clustering was done by using a Euclidian distance and complete clustering method and cut tree to 4 clusters. Each cluster expression was plotted and classified to single or double positive, or double negative of SPARCL1 and FOXJ1 populations.

### Data and code availability

Genome assembly and chromium linked-read sequences were deposited in the DDBJ under accession numbers BLXN01000001–BLXN01022349 and DRA010274. The gene models are available from Figshare under the DOI: 10.6084/m9.figshare.12807032. Single-cell RNA-seq data have been deposited in the DDBJ under the DRR Run accession number DRR496478- DRR496488. Codes used for analysis can be found in https://github.com/wuquan723/Ferret-single-cell-data-from-Matsuzaki-lab. Any information required to obtain and reanalyze the data reported in this paper is available from the lead contact upon request.

## Supporting information

List of Supplementary files

## Acknowledgements

We thank Kazuaki Yamaguchi, Rohab F. Abdelhamid, and Chiharu Tanegashima for sequencing RNA and genomic DNA and all members of the Laboratory of Cell Asymmetry for providing technical support and helpful discussions. We also thank an anonymous reviewer 1 of Review Commons for his/her constructive suggestions, and Fumiya Kusumoto for his continuous encouragement. This work was supported by JSPS KAKENHI [Grant Numbers 18H04003, 17H05779, 19H04791] and RIKEN. RIKEN funds F.M. M.B. was a RIKEN International Program Associate. Q.W. was supported by a JSPS Postdoctoral Fellowship and a RIKEN Special Postdoctoral Researcher Program.

## Author contributions

M.B., Q.W., and F.M. designed experiments. M.K., O.N., Q.W., Y.K., and S.K. constructed new ferret gene models. M.B. and Q.W. performed scRNA-seq experiments and bioinformatics analyses. M.B., Y.T., and Q.W. performed molecular biological analyses. M.B., T.S. [Taeko] and T.S. [Tomomi] performed histological analyses. A.S and Q.W. performed and analyzed Rv NA/protein double-tissue staining. T.S. [Taeko] carried out time-lapse imaging. T.S. [Taeko], Y.T. and H.K. bred the ferrets. M.B., Q.W., A.S. and F.M. wrote the manuscript. All authors read and approved the final manuscript for submission.

## FIGURE LEGENDS

## Supplementary Figure Legends

**Figure S1.**
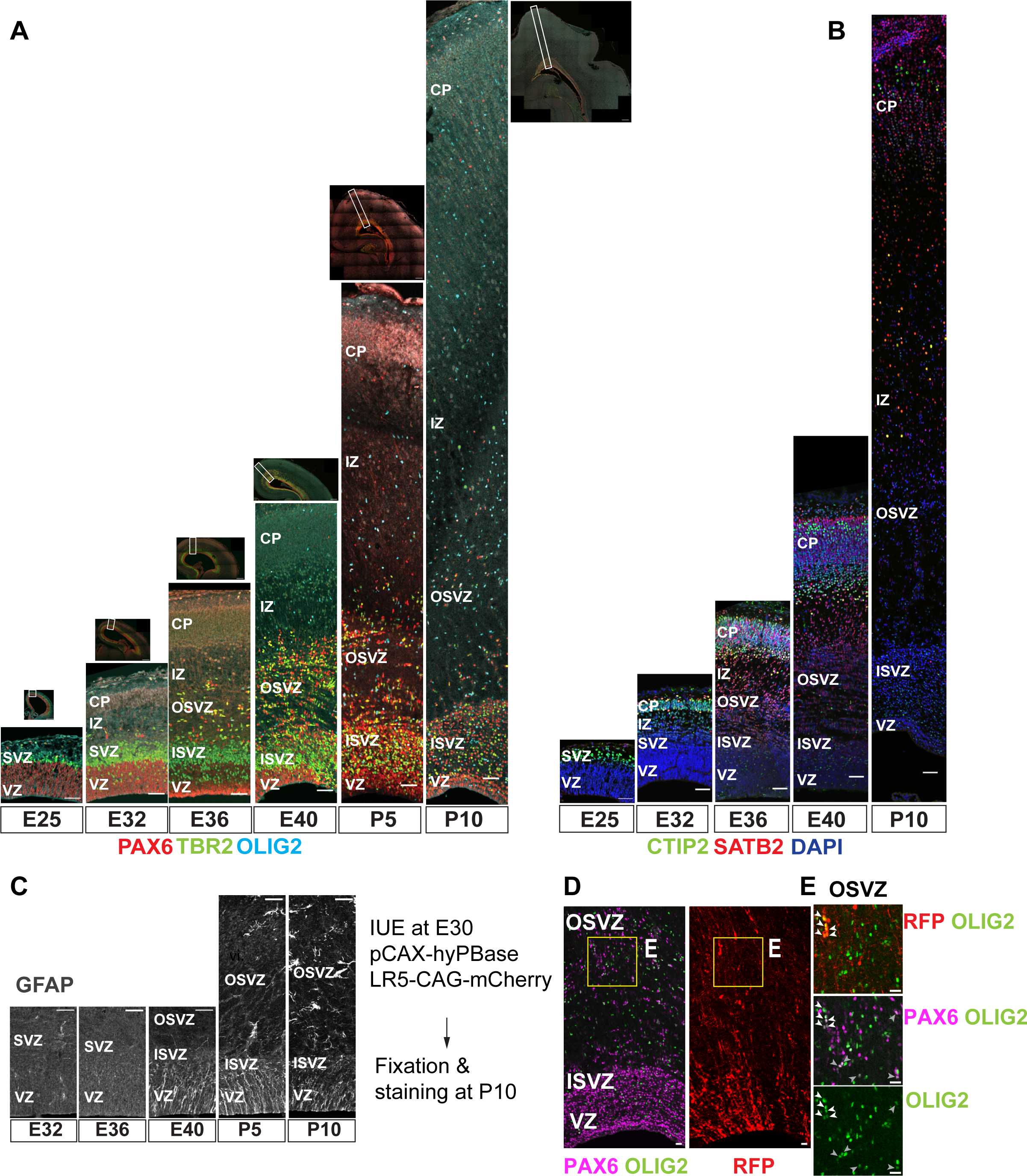
Temporal pattern of neurogenesis and gliogenesis in the cerebral cortex of ferrets. **A** Coronal section of the dorsal cortices immunostained for RG (PAX6, red), IPC (TBR2, green), and OPC (OLIG2, cyan) from early to late neurogenesis at E25, E32, E36, E40, P5, and P10. Scale bars = 100 µm. The position of the image strip at each stage is shown in the corresponding dorsal hemisphere image above the strip. The approximative boundaries of dorsal cortex area used for single-cell RNA sequencing are highlighted with dotty line segments in the dorsal cortex hemisphere above each strip. **B** Immunostaining for CTIP2 (green) and SATB2 (red) at the same developmental stages as in (**A**), showing the onset of DL- (E25 or earlier) and UL-neurogenesis (E32–E34). Scale bars = 100 µm. **C** Immunostaining for GFAP, a marker showing gliogenic potential, in cortical germinal layers at E32, E36, E40, P5, and P10. Gliogenic progenitors emerge around E40, while mature astrocytes with a typical astrocytic morphology appear later at approximately P10 onward in the outside of the germinal layers (Reillo & Borrell, 2012). **D, E** mCherry labeling of the dorsal cortex at E30 showing that a population of OLIG2^+^ oligodendrocyte precursors are generated in the cerebral cortex by P10. **E.** As reported for both mice and humans (Zheng *et al*, 2018; Rash *et al*, 2019b; Huang *et al*, 2020; Kessaris *et al*, 2006). Arrowheads in the cropped images of the left panels (E area in Fig. S1**D**) indicate cells that co-express mCherry, PAX6, and OLIG2 in the OSVZ. Scale bars = 20 µm.

**Figure S2.**
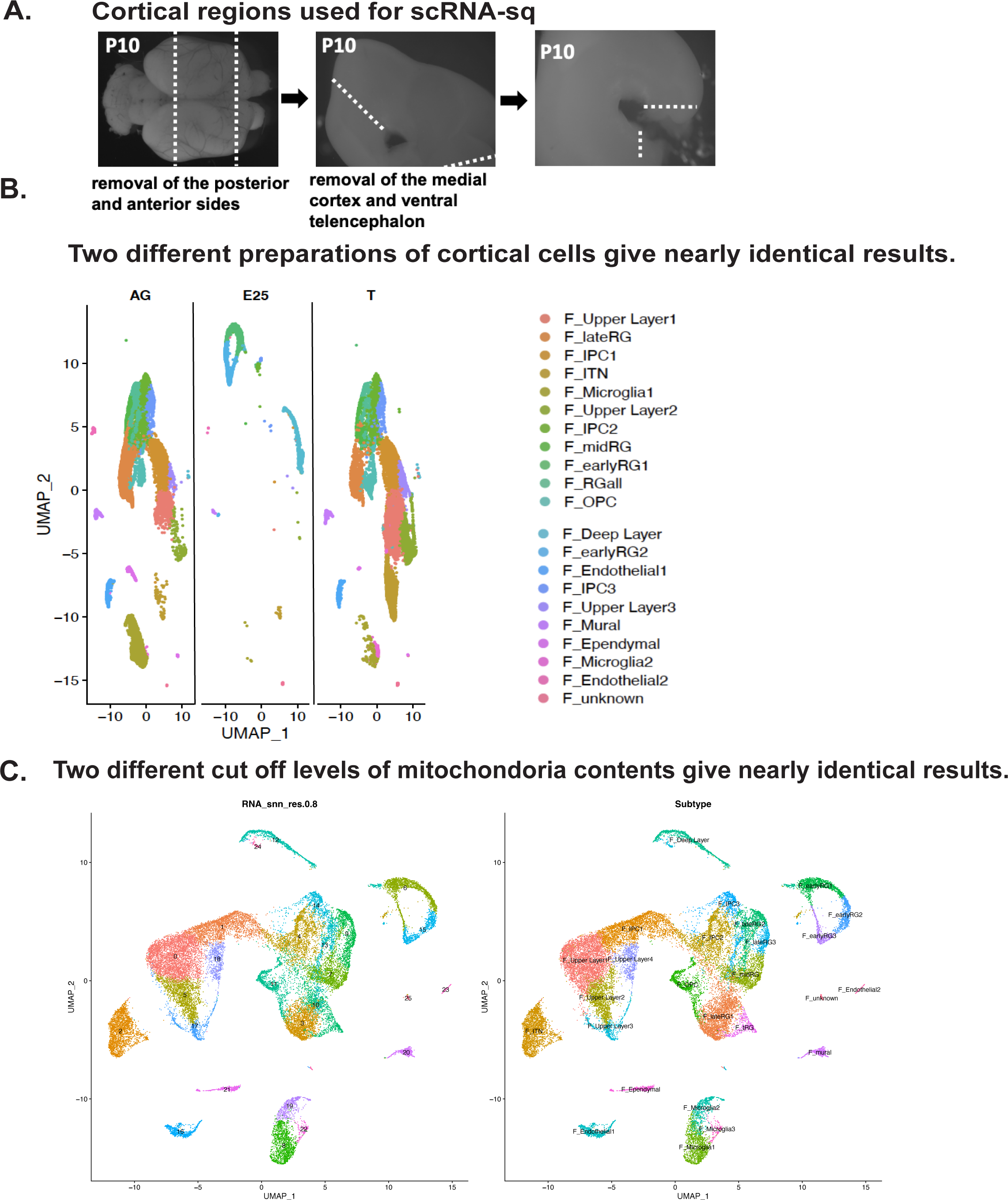
– Quality and reproducibility of ferret single cell transcriptome dataset. **A** The brain region used for the cellular isolation for a scRNA-seq are shown. Dotty lines indicate the approximative boundaries where the cortical slices were cut. **B** Separate UMAP plots for each cell collection series that was prepared by different methods, after merged clustering. Left is the “AG” sample: FACS-based sorting of the neural stem cell fraction labeled with an AzamiGreen (AG)-driven by *HES5* promoter (Ohtsuka et al, 2006). The “T” samples are shown in the right: cells forming the VZ, SVZ, and intermediate zones (IZ) of cerebral cortices after discarding the cortical plate. E25 cells, which were mostly RG, were not collected by cell sorting and put between AG and T in UMAP plots. **C** Effects of cell filtration according to the mitochondrial content. We set the threshold to 10%, resulting in 28,686 cells in our dataset, and then performed the workflow from the normalization step with the same settings that had been applied to our original ferret dataset (Methods). Left is clusters after the 10%-filtered subset on UMAP, and the original clusters in **Fig. 2A** is right. Both conditions gave 26 clusters, and both major cell types and subtypes were conserved after filtering.

**Figure S3.**
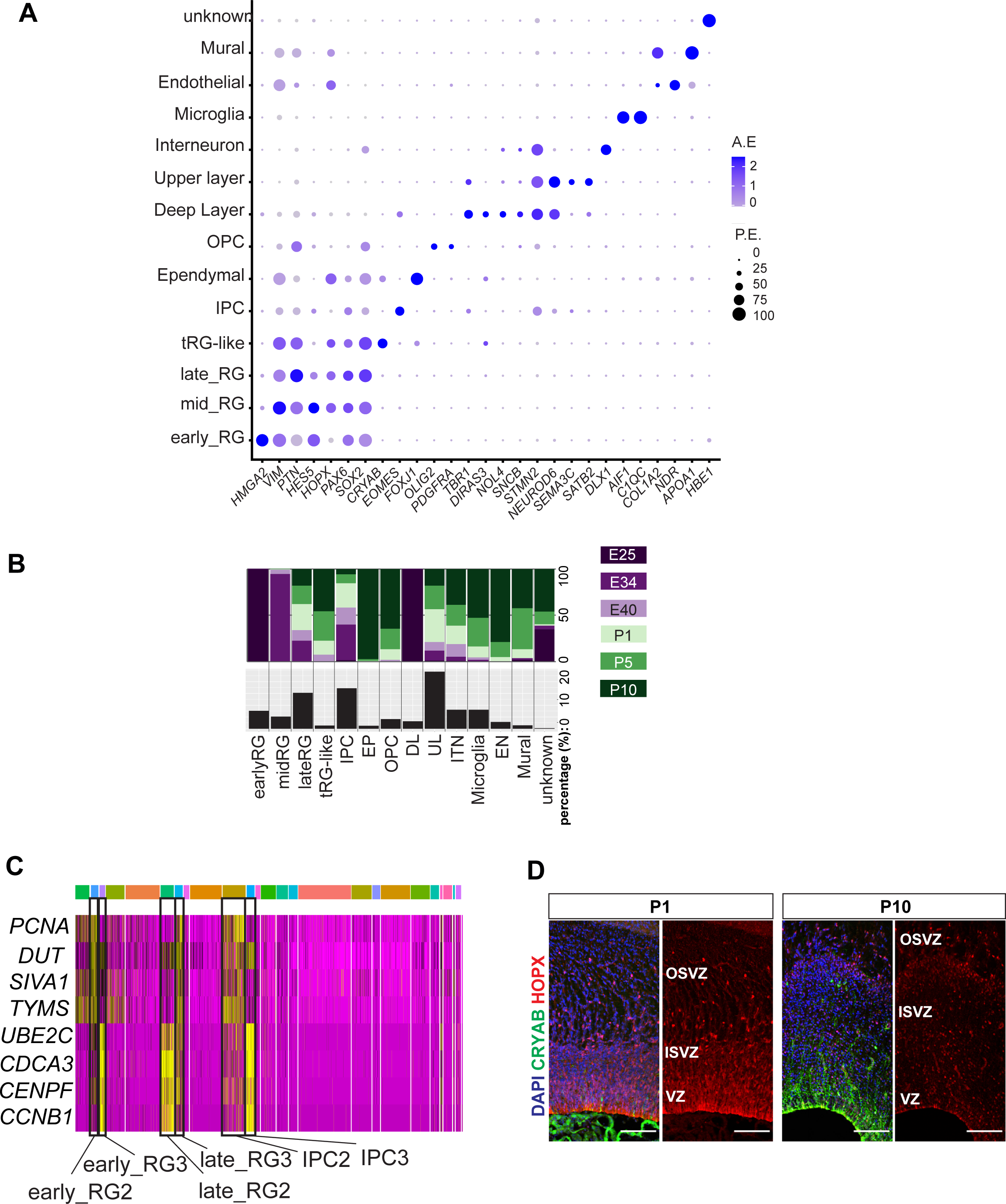
Cell types in developing ferret cerebral cortex. **A.** Dotplot showing average expression and percentage of expression of representative marker genes for individual cell types in the merged dataset. The largest cluster was composed of UL neurons, and their four clusters (**Fig. 2A**) were combined as a single “UL” group. P.E., percentage of expressing cells per cell type; A.E., average expression per cell type. **B.** Percentage of sample collection stages in each cell type (top half) and percentage of each cell type in the merged dataset (bottom half). **C.** Heatmap showing the expression level of cell cycle marker genes based on log fold-change values. *PCNA*, *DUT*, *SIVA1,* and *TYMS* are markers of the S-phase; *UBE2C*, *CDCA4*, *CENPF,* and *CCNB1* are markers of the G2M-phase. Three subclusters among early RG, late RG, and IPC clusters expressed different markers for cell-cycle states. The color bar at the top indicates Seurat clusters matching those in **Fig. 2A**. **D.** Immunostaining for CRYAB^+^ tRG (green) and HOPX^+^ RG (red) together with DAPI on P1 and P10 germinal layers.

**Figure S4.**
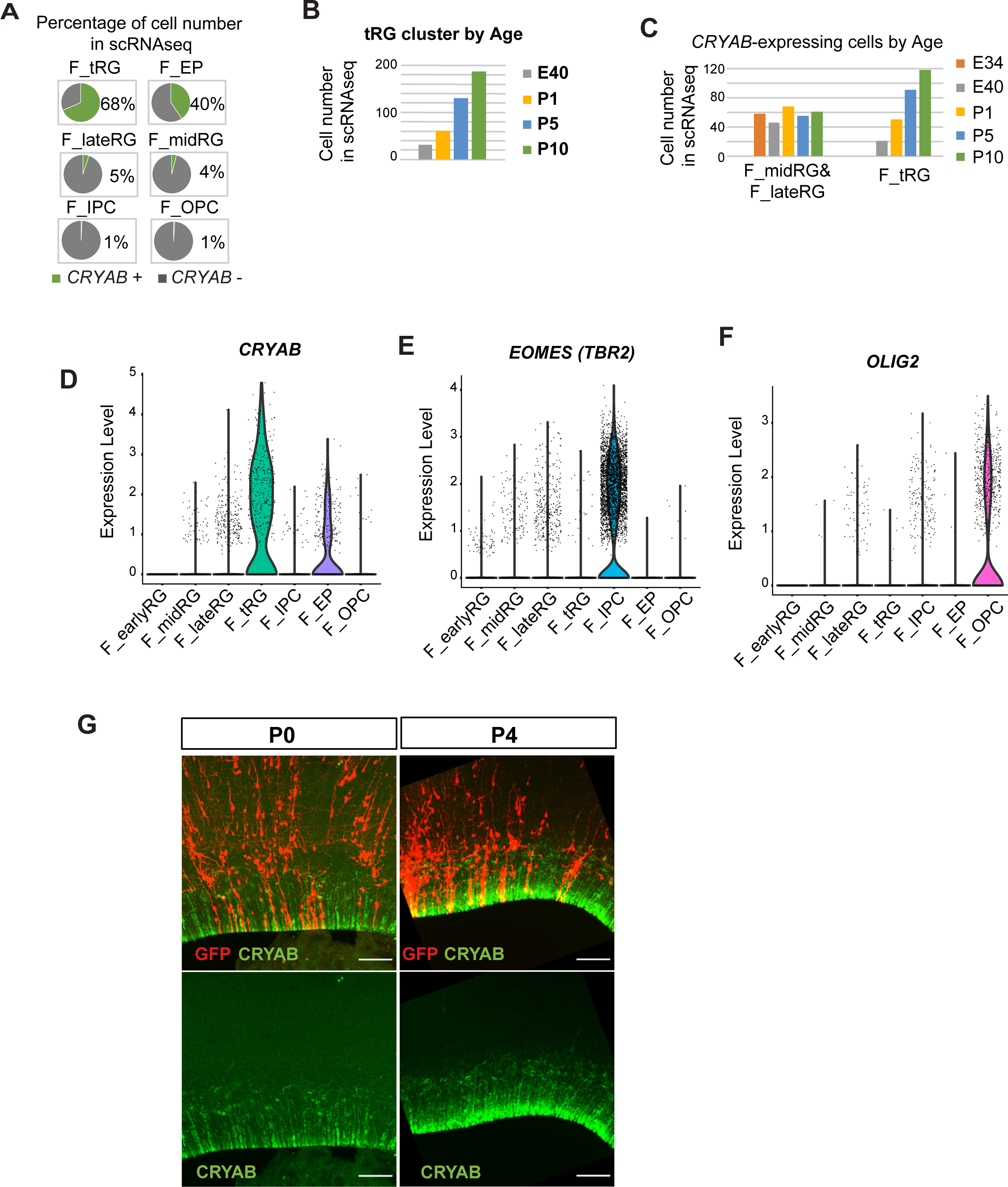
Development of ferret tRG during postnatal cortical development. **A**Pie charts showing the percentage of *CRYAB* expression in the ferret clusters of the merged transcriptome dataset of all stages (**Fig. 2**). Cells were counted as positive if unique molecular identifier counts for *CRYAB* were > 1 using Seurat “WhichCells” function (slot = “counts”). **B** Number of cells in the tRG clustered by age annotated in the ferret transcriptome dataset (**Fig. 2**). **C**. Number of *CRYAB*-expressing cells in RG clusters of the transcriptome dataset (Fig. 2). *CRYAB* expression was undetectable at E25. *CRYAB*-expressing cells in the midRG and lateRG subtypes did not change in all stages (cell numbers: 58, 46, 68, 55, and 61 at E34, E40, P1, P5, and P10, respectively), while the number of *CRYAB*^+^ tRG cells increased after birth (cell numbers: 0, 21, 50, 91, and 118 at E34, E40, P1, P5, and P10, respectively). **D–F.** Violin plots indicating the normalized expression of *CRYAB* (D), *TBR2* or *EOMES* (E), and *OLIG2* (F) in the ferret clusters shown in Fig. 2. **G.** Cellular features of a tRG cell in cortex labelled with GFP at embryonic stages via IUE. Vibratome sections at P0 and P4 (thickness = 200 µm) were immunostained

**Figure S5.**
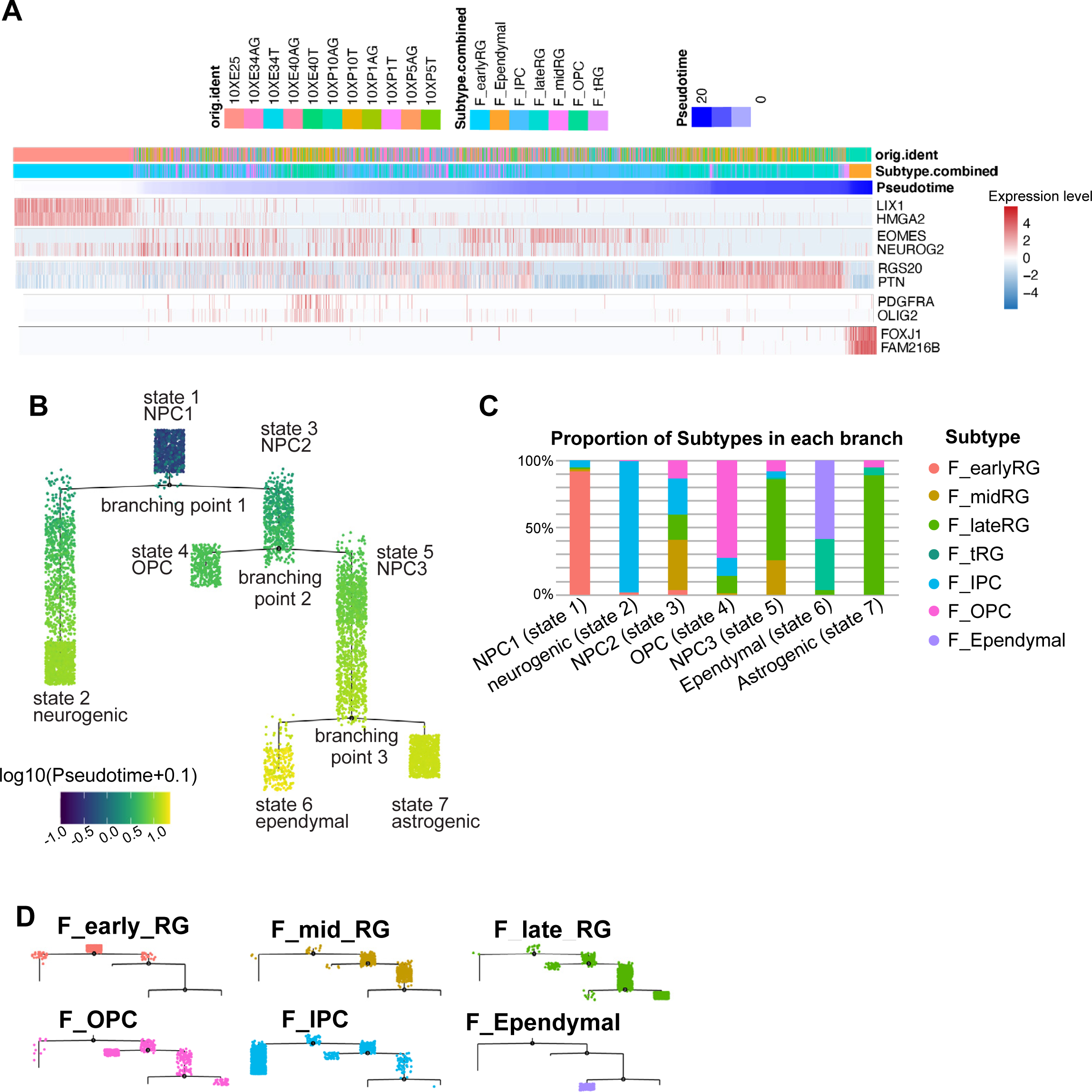
Temporal fates of RG cells identified by pseudo-time trajectory analysis. **A.** Ferret cells used for pseudo-time trajectory analysis were lined up along the stretched pseudo-time axis. Expression levels of markers for typical cell types are also shown along the pseudo-time axis: *LIX1* and *HMGA2* for early RG, *EOMES* and *NEUROG2* for IP, *RGS20*, and *PTN* for late RG, *PDGFR* and *OLIG2* for glial cells, and *FOXJ1* and *FAM216B* for ependymal cells. **B.** Branching trees labelled by pseudo-time score. Color scale indicates log10 (pseudo-time +0.1) values. Each state is named after the major cell type. **C.** The composition of each state is shown by clusters defined in the UMAP plot (**Fig. 2A**). Percentages for clusters were calculated from all cells in each state. **D.** Branching trees split by UMAP clusters (**Fig. 2**).

**Figure S6.**
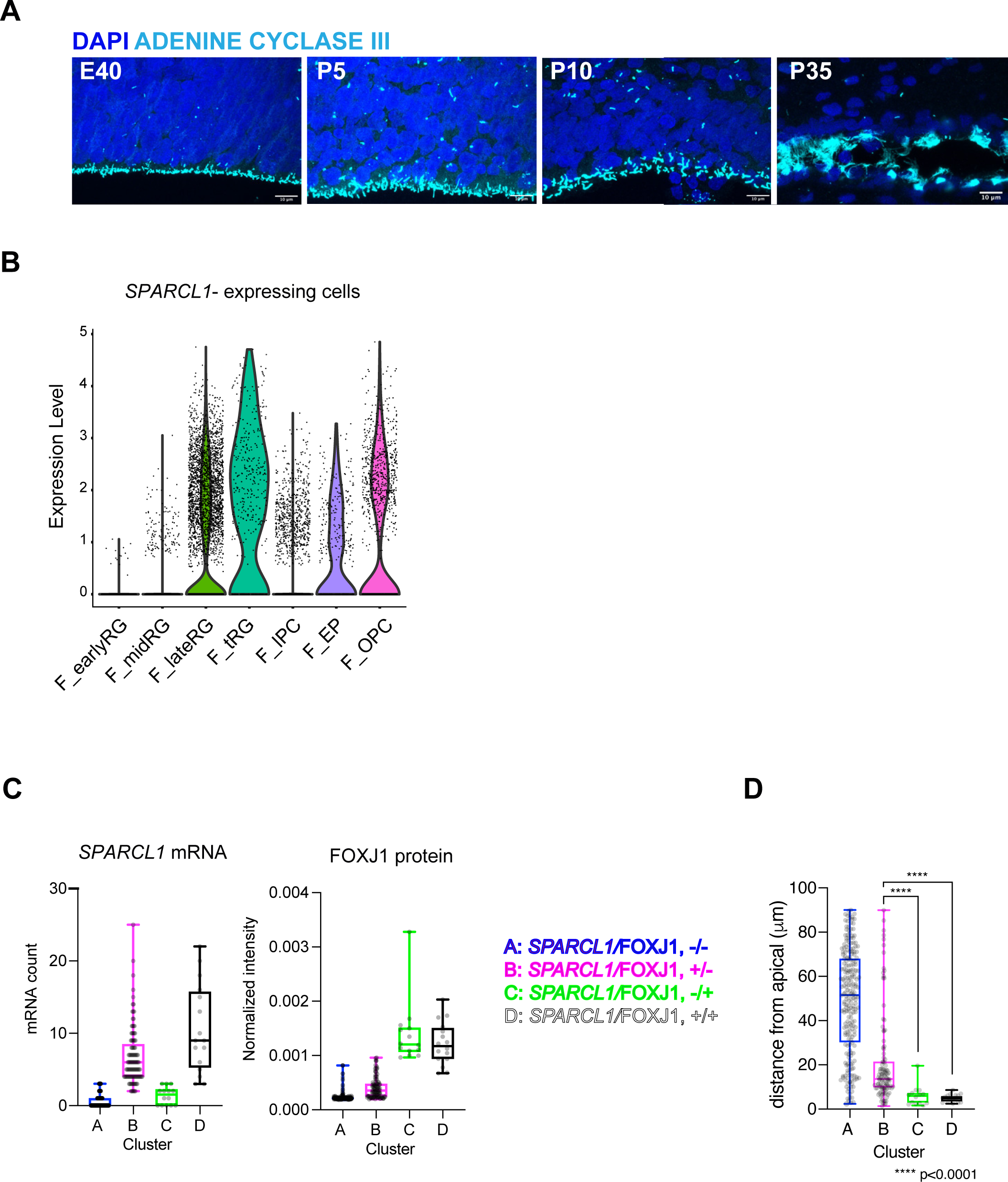
Ferret tRG adopt both ependymal and astrogenic fates in ferrets. **A**. Onset of ciliogenesis in the developing cortex of ferrets, shown via staining for ADENINE CYCLASE III (cyan) along the VZ surface at E40, P5, P10, and P35. Scale bars = 10 µm. **B**. Normalized expression of *SPARCL1*/cell in the major cell types shown in **Fig. 2A**. **C**. Plots showing *SPARCL1* mRNA count (left) and normalized FOXJ1 protein levels (right) in each cell clustered into individual classes (A–D) within the VZ shown in Fig. 4H (Table S4). The box and whisker plots indicate the range (lines) and the upper and lower quartiles with the median (box). D. Distances of each cell (its center of gravity) from the apical surface in the individual clusters **(A–D**), as indicated by the box and whisker plots in (C). Cluster C and D cells are significantly located on the apical surface compared to cluster B cells. Data normality was analyzed by Anderson-Darling test and statical significance was calculated by Kruskal-Wallis test with Dunn’s multiple comparisons test.

**Figure S7.**
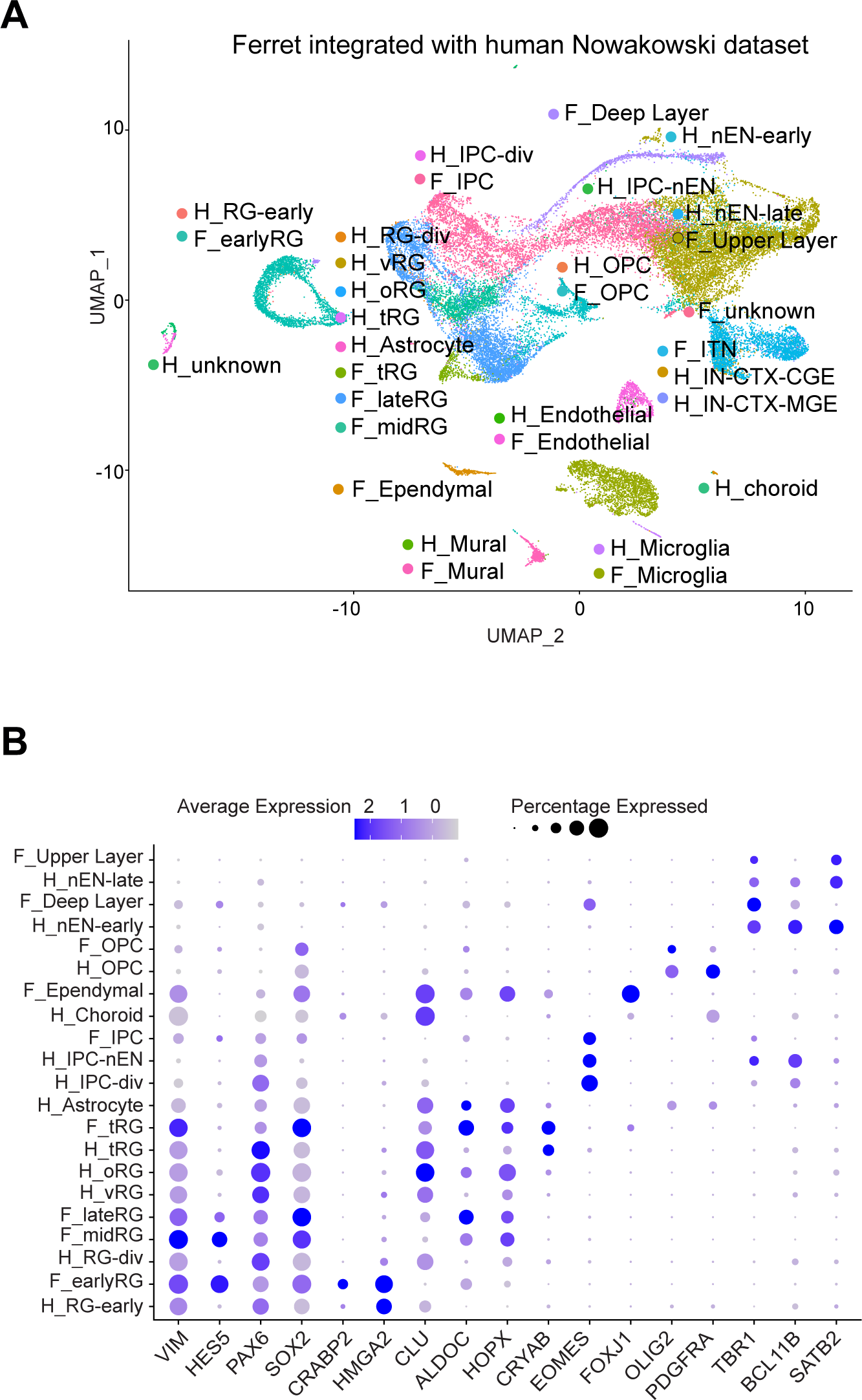
Comparison of molecular identity of RG subtypes between ferret and human. **A.** UMAP visualization of integrated ferret (n = 30,234) and human (n = 2,672) single-cell datasets colored according to the different clusters. The names of clusters from human and ferret cells begin with “H” and “F,” respectively. **B.** Expression patterns of the indicated genes in each cluster. Circle color indicates the expression level of each gene. Circles size indicates the percentage of cells expressing the gene in the indicated cluster.

**Figure S8.**
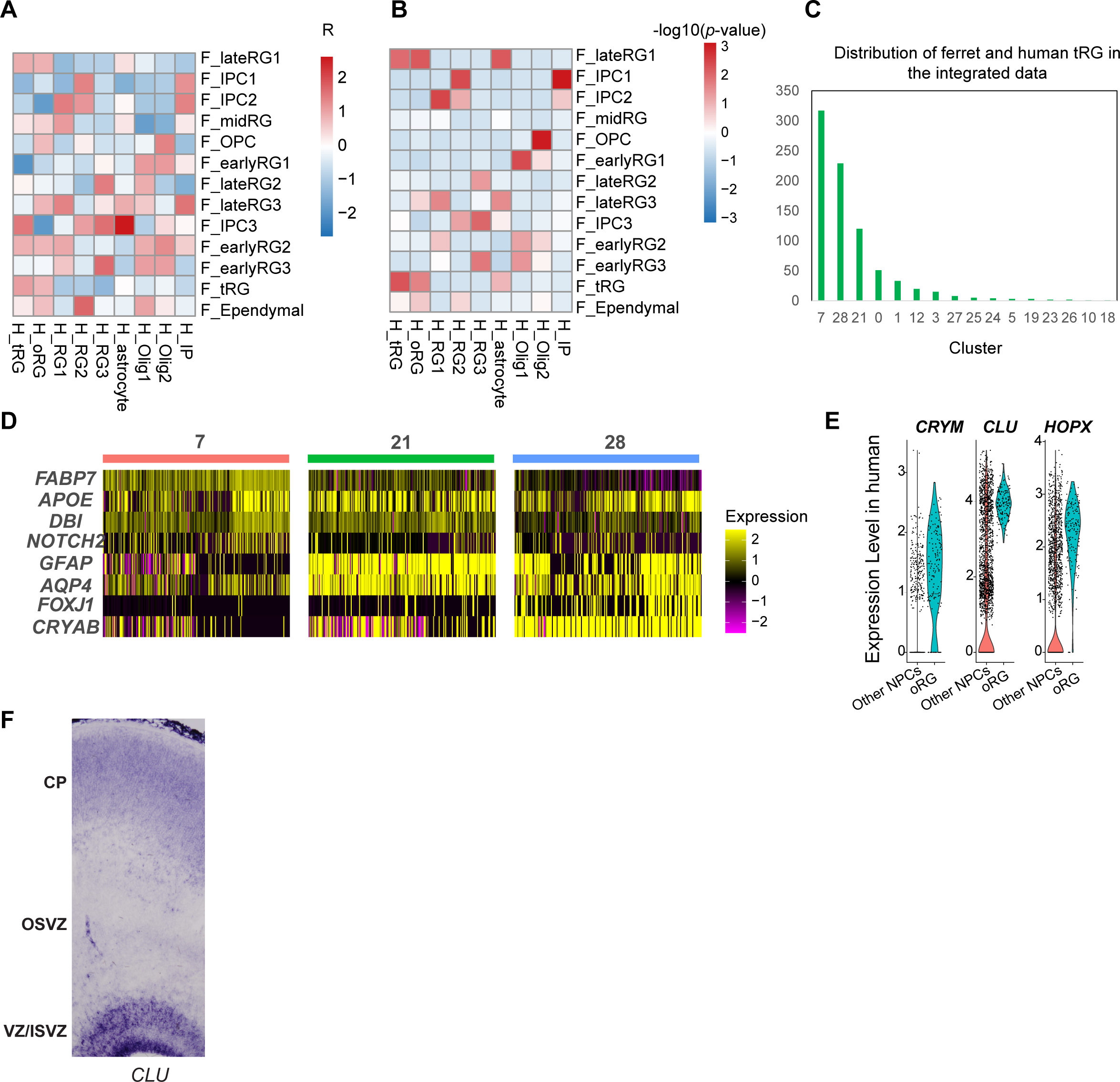
Comparison of the tRG and oRG subtypes between ferrets and humans. **A, B** Matrices of correlations between two arbitrary RG subtypes from humans and ferrets by calculating marker gene scores as described for Fig. 5. (**A**) Correlations and (**B**) *p*-values between pairs in human and ferret clusters are shown. The early RG in ferrets showed the highest similarity with “OLIG1” in human data. This might be because of two reasons; (1) cells only at GW25 without early RG were used for comparison, and (2) cells in “OLIG1” highly expressed genes related with cell proliferation, as the early RG also did (**Table S6**). We did not observe any cell types in human data corresponding to the ferret “mid RG” with the same reason described above. **C** Distribution of human and ferret tRGs in the integrated dataset. The number in the X-axis indicates the cluster number identified by the UMAP in **Fig. 7C**. **D** Expression levels of the subtype maker genes in the three tRG-enriched clusters (7, 21, and 28). Marker enrichment is consistent with the presumptive identification of tRG subtypes according to pseudotime trajectory analysis. **E** Expression patterns of *CRYM*, *CLU*, and *HOPX* in oRG cells and other NPCs in humans. **F** *In situ* hybridization of *CLU* at the ferret cortex showed its expression in all VZ, ISVZ, and OSVZ at P5.

**Figure S9.**
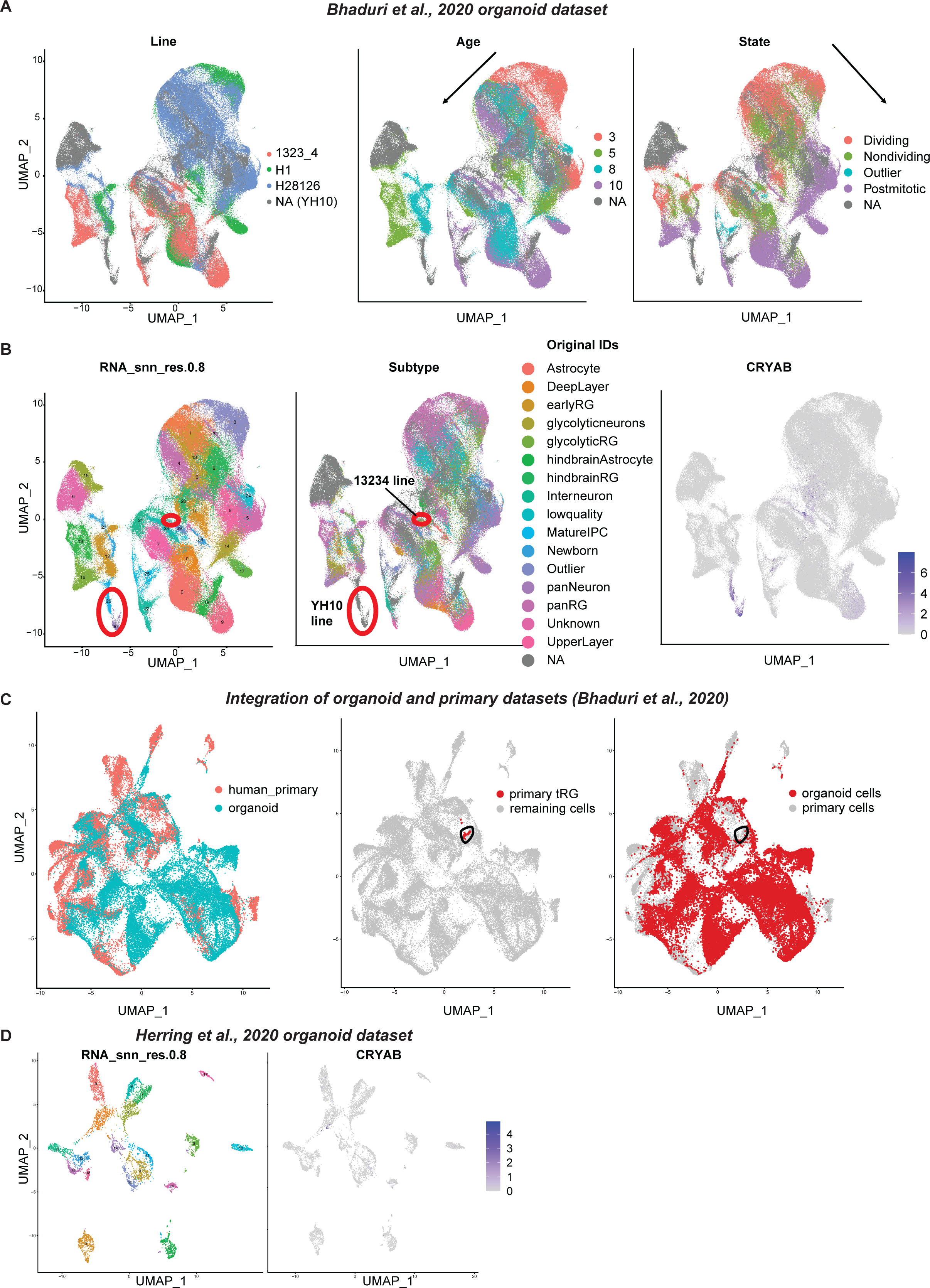
The absence of a tRG cluster in human organoid datasets. **A**. UMAP visualization of human organoid-driven cells colored by the original annotations for the cell line (left), age (middle) or state (right). Human organoid dataset from Bhaduri et al., 2020 was used. Bhaduri dataset contained organoids generated from 4 different lines, which showed a variability in terms of cell distribution on UMAP while overall temporal and differentiation axes were recapitulated. **B**. UMAP visualization of cells by clusters (left), original cluster annotations (middle), or by the normalized expression level of *CRYAB* (right). Red circles on UMAP highlight the group of cells expressing *CRYAB*. *CRYAB* expression was restricted to two out of four cell lines. CRYAB was expressed in clusters 26 and 30 enriched in the YH10 line, and cluster 29 enriched in the13234 cell line. **C**. UMAP visualization of integrated primary and organoid datasets from Bhaduri et al., 2020. Cells are colored by the originated dataset (left), primary tRG cluster (middle) or organoid dataset (right). Black circles were used to indicate the overlap of the region where human primary tRG was found. Human tRG did not overlap with organoid cells. We found that CRYAB-expressing organoid clusters 26 and 30 overlapped with “oRG/astrocyte” clusters of primary tissues. **D**. UMAP visualization of cells derived from Herring et al., 2022 human organoid dataset, colored by clusters (left) or by the normalized expression level of *CRYAB* (right).

