## Supplementary material for "Truncated radial glia as a common precursor in the late corticogenesis of gyrencephalic mammals": List of Supplementary files

Table S1: Quality control, metadata, and cluster information, related to Figure 1.

Table S2<-S3: Pseudotime trajectory analysis, related to Figure 4.

Table S3 (new): Expression of *SPARCL1* and *FOXJ1* mRNAs in ferret tRG cells, related to Figure 5.

Table S4 (new): Distances of P10 cells with *SPARCL1* mRNAs ± ,FOXJ1 protein± from the apical surface,

related to Figure 5.

Table S5<-S2: Human-ferret integration; metadata, cluster markers, related to Figure 6.

Table S4->S6: Cluster markers in integrated dataset, related to Figure 7.

Table S5->S7: Extraction of total RNA from different tissues, used for the gene model reconstruction
